# *Giardia intestinalis* reshapes mucosal immunity toward a Type 2 response that attenuates inflammatory bowel-like diseases

**DOI:** 10.1101/2024.03.02.583119

**Authors:** Aline Sardinha-Silva, Pedro H. Gazzinelli-Guimaraes, Oluwaremilekun G. Ajakaye, Tiago R. Ferreira, Eliza V.C. Alves-Ferreira, Erick T. Tjhin, Beth Gregg, Marc Y. Fink, Camila H. Coelho, Steven M. Singer, Michael E. Grigg

**Author notes:** Current address: Center for Drug Evaluation and Research, US Food and Drug Administration, USA. Corresponding author: Aline Sardinha-Silva and Michael E. Grigg. Molecular Parasitology Section, Laboratory of Parasitic Diseases, National Institute of Allergy and Infectious Diseases, National Institutes of Health, Bethesda, MD 20892, United States. and.

## Abstract

Diarrheal diseases are the second leading cause of death in children worldwide. Epidemiological studies show that co-infection with *Giardia intestinalis* decreases the severity of diarrhea. Here, we show that *Giardia* is highly prevalent in the stools of asymptomatic school-aged children. It orchestrates a Th2 mucosal immune response, characterized by increased antigen-specific Th2 cells, IL-25, Type 2-associated cytokines, and goblet cell hyperplasia. *Giardia* infection expanded IL-10-producing Th2 and GATA3^+^ Treg cells that promoted chronic carriage, parasite transmission, and conferred protection against *Toxoplasma gondii*-induced lethal ileitis and DSS-driven colitis by downregulating proinflammatory cytokines, decreasing Th1/Th17 cell frequency, and preventing collateral tissue damage. Protection was dependent on STAT6 signaling, as *Giardia*-infected STAT6^-/-^ mice no longer regulated intestinal bystander inflammation. Our findings demonstrate that *Giardia* infection reshapes mucosal immunity toward a Type 2 response, which confers a mutualistic protection against inflammatory disease processes and identifies a critical role for protists in regulating mucosal defenses.

## INTRODUCTION

The gastrointestinal tract is the largest mucosal surface in all mammalian systems, which harbors a diverse community of commensals; but it is also a target for a multitude of other pathogenic organisms, including viruses, bacteria, helminths, and protozoan parasites, that can cause severe enteric infections (Sardinha-Silva et al., 2022). Therefore, intestinal mucosal immunity plays a critical role by providing the first line of defense against pathogenic organisms, but is also fundamental to promote tolerogenic immune responses to maintain tissue homeostasis (McGhee and Fujihashi, 2012).

Remarkably, diarrheal disease and enteric infections caused by a variety of pathogens constitute the second leading cause of death globally in children (Collaborators, 2018; WHO, 2017). This is especially prevalent in low– and middle-income countries characterized by poor sanitation and a relative lack of adequate water supplies, which contribute significantly to the prevalence of gastrointestinal pathogens and often lead to co-infections. Among these pathogens, *Giardia intestinalis*, a gastrointestinal protozoan parasite, can reach a prevalence of 20-30% among children in low– and middle-income countries (Dixon, 2021).

Establishment of *Giardia* in the small intestine induces a robust Th17 immune response, which restricts *Giardia* infection and is thought to protect the host (Dann et al., 2015; Dreesen et al., 2014). Because Th1 and Th2 immunity were not associated with protection against *Giardia* infection in mice (Singer and Nash, 2000), the induction and regulation of these responses during intestinal infection are not well defined. Experimental *Giardia* infection in humans resulted in diarrhea, leading to the classification of this protist as an enteric pathogen (Nash et al., 1987).

Further, previous work showed that *Giardia* infection is associated with increased intestinal permeability (Muller and von Allmen, 2005; Troeger et al., 2007), microbial translocation (Chen et al., 2013), and intestinal pathology that is mediated by CD8^+^ T cell activation (Keselman et al., 2016; Scott et al., 2004).

However, despite such responses, epidemiological studies indicate that children co-infected with *Giardia intestinalis* alongside other enteric pathogens (mostly *Cryptosporidium*, *Salmonella*, and rotaviruses) exhibit a reduced risk of developing severe life-threatening diarrhea compared to those with single-pathogen infections, highlighting *Giardia*’s role in preventing severe outcomes (Bhavnani et al., 2012; Cotton et al., 2014b; Muhsen et al., 2014; Oberhelman et al., 2001; Veenemans et al., 2011; Wang et al., 2013). In this scenario, *Giardia* is considered a pathobiont, but the underlying mechanism of protection remains enigmatic and controversial. Persistent, largely asymptomatic *Giardia* infection has been reported, and infection can last for months (Das et al., 2023; Rogawski et al., 2017), suggesting that in some hosts, it may exist as a constituent member of the host commensal microbiome.

In this study we used a murine model of *Giardia intestinalis* infection to comprehensively investigate how the parasite influences the initiation of mucosal immunity at single-cell resolution. We show that *Giardia* infection triggers an early antigen-specific Type 2 immune response in the intestine, marked by elevated levels of IL-4 and IL-13, along with an increase in effector Th2 cells (IL-13 producing FoxP3^-^GATA3^+^CD4^+^ T cells) in the small intestine lamina propria (SI LP). This response was associated with goblet cell hyperplasia, increased mucus secretion, and higher IL-25 levels. Furthermore, we identified an expansion of IL-10-producing CD4^+^ Th2 and GATA3^+^ T regulatory (Treg) cells in the intestine, which we established are essential for parasite persistence. Specifically, depleting *Giardia*-induced type-2 immunity in STAT6^-/-^ mice led to a significant reduction of IL-10 levels and a remarkable switch to Th1/Th17 immune responses, increasing inflammation in the small intestine that resulted in the rapid clearance of *Giardia* parasites. Moreover, experiments involving inflammatory bowel-like disease models including a Th1/Th17 colitis driven by dextran sodium sulfate (DSS) and a Th1-driven/IFN-gamma-mediated acute ileitis induced by *Toxoplasma gondii*, an intestinal protozoan parasite that causes a Crohn’s disease-like enteritis, demonstrated *Giardia*’s anti-inflammatory potential, by suppressing IFN-γ producing T-bet^+^ Th1 cells during *T. gondii* co-infection; and dramatically reducing IL-6 levels and pathology in the DSS-induced colitis model. Lastly, we showed that these protective mechanisms were dependent on *Giardia*-driven Th2 immunity and STAT6 signaling in the intestine, as *Giardia*-infected STAT6^-/-^ mice were no longer able to regulate bystander intestinal inflammation. These results underscore a previously unrecognized mutualistic interaction between the host and a commensal protist that may critically impact the course and progression of inflammatory bowel-like diseases (IBDs) such as Crohn’s disease (CD) and Ulcerative colitis (UC).

## RESULTS

### High frequency asymptomatic carriage of *Giardia intestinalis* in school-aged children in Africa

Giardiasis is one of the most prevalent enteric diseases associated with stunting in children under 2 years of age (Rogawski et al., 2018), with prevalence estimates ranging from 2% to in excess of 30% from high and low-middle income countries, respectively (Dixon, 2021). More recently referred to as a commensal pathogen (Sardinha-Silva *et al*., 2022), the degree to which *Giardia* persists as a natural member of the host gut microbiome has not been systematically studied. To investigate the prevalence of asymptomatic carriage of *Giardia* infection in susceptible school-aged children, we screened 664 fecal samples collected across 6 health districts along a longitudinal transect from the North to the South of Nigeria that comprised vastly different ecotypes and was a mix of rural and semi-urban sites (Supp.Fig.1 and Table 1). Quantitative PCR (qPCR) results using genomic DNA extract from fecal samples indicated that the incidence of infection was high in all states, ranging from a low of 55% in Plateau to a high of 86% in Enugu (Table 1). When assessed for infection that was associated with symptomatic disease (abdominal cramps, nausea, fever, diarrhea), 4 districts (Benue, Enugu, Jigawa, Kano) reported very few cases of symptomatic disease, ranging from zero to 9%, relative to Plateau and Cross River, where *Giardia* infection was associated with symptoms in 25 and 80 % of children, respectively (Table 2). The factors that are regulating the different levels of pathogenicity between health districts (*i.e.,* asymptomatic vs. symptomatic carriage) are unclear, and are being investigated further. However, it is clear from this study, that the vast majority of school-aged infections (83%; Table 2) resulted in no obvious disease sequelae, supporting the distinction that asymptomatic carriage of *Giardia intestinalis* is common, widespread, and underappreciated with respect to human infection.

**Table 1:** *Giardia intestinalis* prevalence in children ≤ 10 years-old in Nigeria.

| ID | Locations | *n* | Non-infected, % (*n*) | Infected, % (*n*) |
| --- | --- | --- | --- | --- |
| 1 | Benue | 94 | 19.1 (18) | 80.9 (76) |
| 2 | Cross River | 96 | 21.9 (21) | 78.1 (75) |
| 3 | Enugu | 99 | 14.1 (14) | 85.9 (85) |
| 4 | Jigawa | 92 | 15.2 (14) | 84.8 (78) |
| 5 | Kano | 196 | 20.9 (41) | 79.1 (155) |
| 6 | Plateau | 87 | 44.8 (39) | 55.2 (48) |
| **Total prevalence** |  | **664** | **22.1 (147)** | **77.9 (517)** |

**Table 2:** Asymptomatic vs. Symptomatic infected children ≤10 years-old in Nigeria.

| ID | Locations | *n* | Asymptomatic, % (*n*) | Symptomatic, % (*n*) |
| --- | --- | --- | --- | --- |
| 1 | Benue | 76 | 97.4 (74) | 2.6 (2) |
| 2 | Cross River | 75 | 20.0 (15) | 80.2 (60) |
| 3 | Enugu | 85 | 90.6 (77) | 9.4 (8) |
| 4 | Jigawa | 78 | 100.0 (78) | 0.0 (0) |
| 5 | Kano | 155 | 94.8 (147) | 5.2 (8) |
| 6 | Plateau | 48 | 75.0 (36) | 25.0 (12) |
| **Total positive cases** |  | **517** | **82.6 (427)** | **17.4 (90)** |

### *Giardia* induces an early antigen-specific type-2 immune response in the intestine of infected mice

The impact of asymptomatic carriage of *Giardia* on the host has been practically overlooked, and how it shapes the intestinal immune landscape has not been systematically investigated. To better understand the mucosal immune response induced during intestinal colonization of *Giardia*, we utilized a mouse model of giardiasis by orally infecting C57BL/6 WT mice with 1 x 10^6^ trophozoites of the GS strain (Fig. 1A) and we performed a temporal analysis of cytokine profiling in the small intestine tissue homogenate (Supp.Fig.2). We found that *Giardia* induces increased secretion of Type 2-associated cytokines, such as IL-4 and IL-13, peaking at five to seven days post-infection (Supp.Fig.2B-C), along with IL-17A, which peaked between five– and nine-days post-infection (Supp.Fig.2D). Based on these findings, we next performed a comprehensive characterization of the gut mucosal immune response during *Giardia* infection, seven days post-infection, in both the small and large intestines. Although we did not observe any differences in the numbers of effector IFN-γ producing FoxP3^-^Tbet^+^ Th1 cells, our immunophenotypic analysis (gating strategy in Supp.Fig.3A) revealed that *Giardia* infection induced a pronounced increase in the number of effector IL-13 producing FoxP3^-^GATA3^+^ Th2 cells and IL-17A producing FoxP3^-^RORγt^+^ Th17 cells in the small intestine (Fig.1B). These enriched Th2 and Th17 responses were associated with a marked increase in the small intestine levels of IL-4, IL-13 and IL-17A (Fig.1C). Notably, this mixed cytokine signature was observed to be specific to *Giardia* antigens, as SI LP cells from *Giardia*-infected mice stimulated *in vitro* with *Giardia* soluble antigens induced a prominent increase of antigen-driven IL-4, IL-13 and IL-17 levels, when compared to stimulated cells from naïve mice, or unstimulated cells from both groups (Fig.1D). Furthermore, *Giardia* infection was found to induce goblet cell hyperplasia (Fig.1E), particularly evident in the small intestine, concomitant with elevated levels of IL-25 within the lamina propria tissue (Fig.1F). As expected, our data revealed that *Giardia* trophozoites were only present in the small intestine and were not detected in the colon (Fig.1H). However, the parasite-induced Th2 immune response was observed primarily within the large intestine and was associated with an elevated frequency and absolute number of FoxP3^-^GATA3^+^ Th2 cells, as well as increased IL-4 levels in the colon of *Giardia*-infected mice, in comparison with the naïve group (Fig.1G). These findings unravel the complex site-specific immunological and cellular responses triggered by *Giardia* infection in the small and large intestines, emphasizing the distinct nature of host-pathogen interactions in different segments of the gastrointestinal tract.

**Figure 1.**
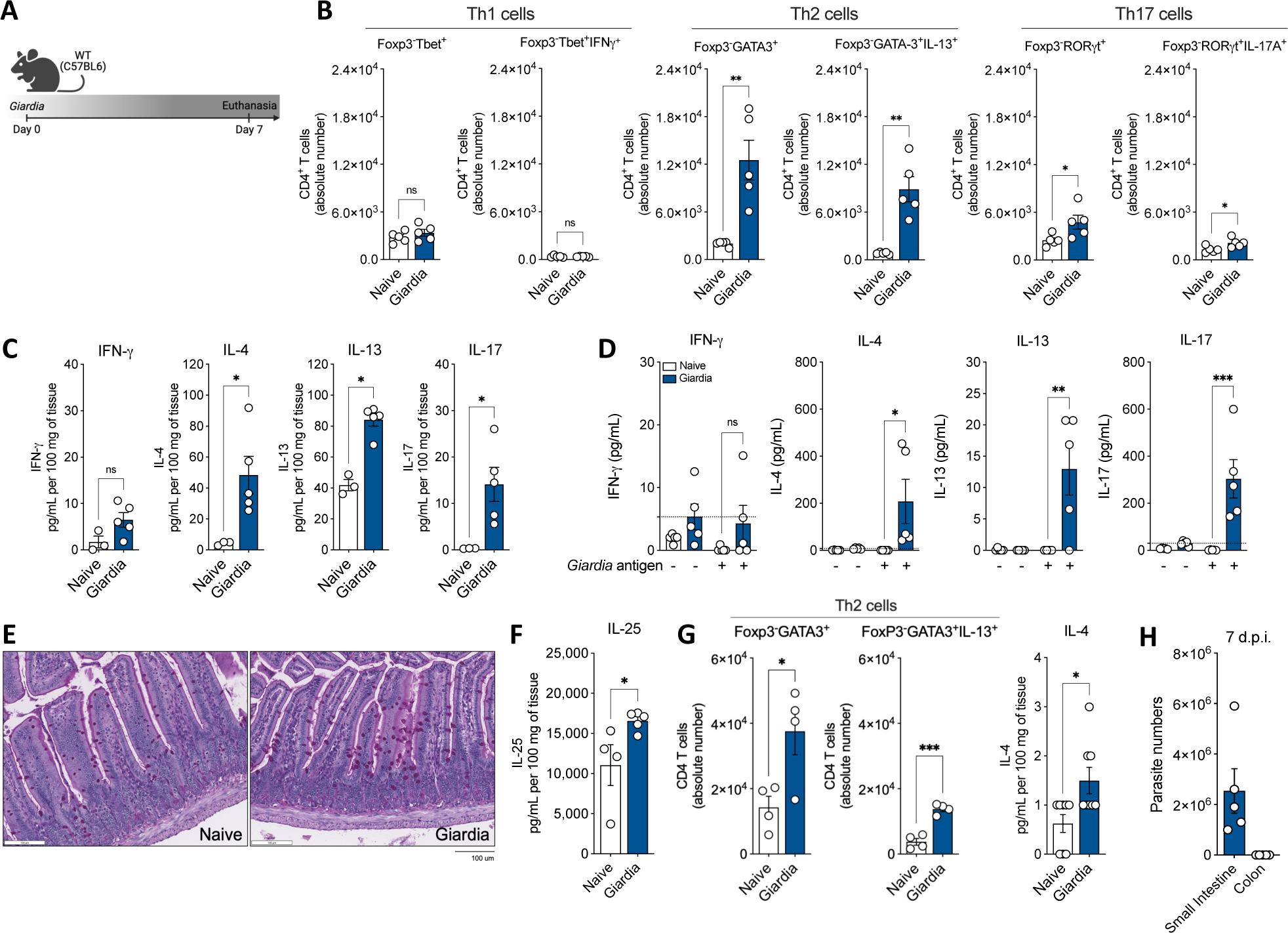
*Giardia*-infected mice induce a Th2 immune response in the lamina propria at the small and large intestines. (A) Experimental schematic. Female C57BL/6 mice were perorally infected with 1×10^6^ *Giardia* GS/M strain trophozoites. (B) Scatter plot graphs indicating the absolute number of Th1 (Foxp3^-^Tbet^+^IFN-γ^+^), Th2 (Foxp3^-^GATA3^+^IL-13^+^), and Th17 (Foxp3^-^ RORγt^+^IL-17^+^) cells in the small intestine lamina propria of *Giardia*-infected mice (7 d.p.i.). Gated on Live CD45^+^TCR3^+^CD4^+^. (C) IFN-γ, IL-4, IL-13, and IL-17 levels in the small intestine tissue homogenate of *Giardia*-infected mice (7 d.p.i.) measured by Luminex. (D) IFN-γ, IL-4, IL-13, and IL-17 levels produced by small intestine isolated lamina propria cells after 24 hours of *in vitro* stimulation with *Giardia* soluble antigens (10 μg/mL). (E) Representative image of the jejunum from naïve or *Giardia*-infected mice stained with Periodic Acid Schiff (PAS) for mucus production by Goblet cells (7 d.p.i.). Scale bars represent 100 μm. (F) IL-25 levels in the small intestine tissue homogenate of *Giardia*-infected mice (7 d.p.i.) measured by ELISA. (G) Scatter plot graphs indicating the frequency and the absolute number of Th2 cells (Foxp3^-^GATA3^+^) in the colonic lamina propria of *Giardia*-infected mice (7 d.p.i.) (gated on Live CD45^+^TCR3^+^CD4^+^), and IL-4 levels in the colon (7 d.p.i.). (H) *Giardia* burden in the small intestine (duodenum) and colon of C57BL/6 infected mice (7 d.p.i.). Data are represented as mean ± SEM and significance was calculated with non-parametric Mann-Whitney test. *p≤0.05, **p≤0.01, ***p≤0.001. Data are representative of four (B-C) or two (D-H) independent experiments.

### Effector Th2 cells and GATA3^+^ Tregs are the major sources of *Giardia*-driven IL-10 in the small intestine

In addition to the dominant Type 2 and IL-17 cytokines, *Giardia* infection was also associated with an increased production of the regulatory cytokine IL-10 in the small intestine (Fig.2A and Supp.Fig.2E), but not in the colon (Fig.2B). As observed for the other relevant cytokines elicited by *Giardia* (IL-4, IL-13 and IL-17), IL-10 secretion in the small intestine of infected mice was induced by *Giardia* antigen stimulation *in vitro* (Fig.2C). However, the frequency and absolute number of IL-10-producing FoxP3^+^GATA3^-^ Treg cells were similar between infected and naïve mice, suggesting a different source of *Giardia*-driven IL-10 than the canonical Treg (Fig.2D). To investigate the source of IL-10 during *Giardia* infection, we orally infected IL-10 GFP reporter mice with 1 x 10^6^ *Giardia* trophozoites and profiled the IL-10 producing cells in the small intestine lamina propria at 7 days-post infection (d.p.i.) (Fig.2E). Flow cytometric analysis demonstrated that in naïve mice, both CD4^+^ T cells and macrophages together represent the major sources of IL-10. Nevertheless, the IL-10-producing CD4^+^ T cells within the lamina propria undergo substantial expansion in response to *Giardia* infection (Fig.2F). Remarkably, the deconvolution of different subpopulations of CD4^+^ T cells demonstrated that the frequencies and absolute numbers of IL-10-producing FoxP3^-^GATA3^+^ Th2 cells, and IL-10-producing GATA3^+^ Tregs were dramatically increased in the small intestine of *Giardia*-infected mice when compared with the naive tissue (Fig.2G). Taken together, our data demonstrate that *Giardia* infection triggers elevated levels of IL-10 at the site of infection, as well as expansion of IL-10-producing Th2 and GATA3^+^ Treg cells, which serve as the main source of this anti-inflammatory cytokine.

**Figure 2.**
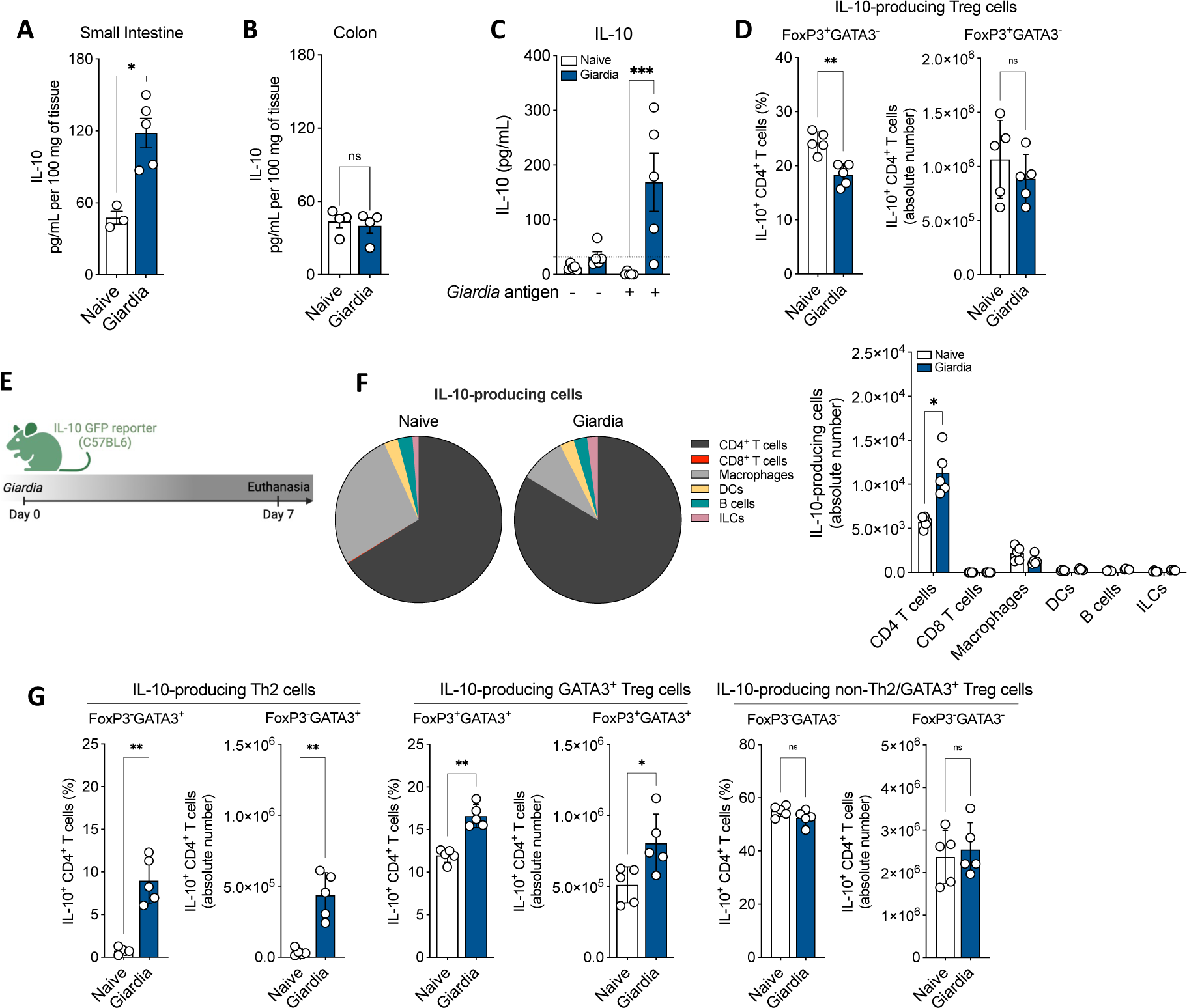
*Giardia* induces secretion of IL-10 and expansion of IL-10-producing Th2 and GATA3^+^ Treg cells in the small intestine. IL-10 levels in the (A) small intestine and (B) colon of *Giardia*-infected mice (7 d.p.i.) measure by Luminex. (C) IL-10 levels produced by small intestine isolated lamina propria cells after 24 hours of *in vitro* stimulation with *Giardia* soluble antigens (10 μg/mL). (D) Scatter plot graphs indicating the frequency and the absolute number of IL-10-producing Treg cells (Foxp3^+^GATA3^-^IL-10^+^) in the small intestine lamina propria of *Giardia*-infected mice (7 d.p.i.) (gated on Live CD45^+^TCR3^+^CD4^+^). (E) Experimental schematic. Female C57BL/6 IL-10 GFP reporter mice were perorally infected with 1×10^6^ *Giardia* GS/M strain trophozoites. (F) Frequency and absolute number of IL-10-producing cells in the small intestine lamina propria of *Giardia*-infected mice (7 d.p.i.). Gated on: CD4^+^ T cells (Live CD45^+^IL-10^+^TCR3^+^CD19^-^CD4^+^), CD8^+^ T cells (Live CD45^+^IL-10^+^TCR3^+^CD19^-^CD8^+^), Macrophages (Live CD45^+^IL-10^+^TCR3^-^CD19^-^CD11b^+^CD64^+^), Dendritic Cells (DCs) (Live CD45^+^IL-10^+^TCR3^-^CD19^-^CD11c^+^MHCII^+^), B cells (Live CD45^+^IL-10^+^TCR3^-^CD19^+^), and Innate Lymphoid Cells (ILCs) (Live CD45^+^IL-10^+^TCR3^-^CD19^-^CD90.2^+^). (G) Scatter plot graphs indicating the frequency and the absolute number of IL-10-producing Th2 (Foxp3^-^GATA3^+^), GATA3^+^ Treg (Foxp3^+^GATA3^+^), and non-Th2/GATA3^+^ Treg (Foxp3^-^GATA3^-^) cells in the small intestine lamina propria of *Giardia*-infected mice (7 d.p.i.) (gated on Live CD45^+^TCR3^+^CD4^+^). Data are represented as mean ± SEM and significance was calculated with non-parametric Mann-Whitney test. *p≤0.05, **p≤0.01, ***p≤0.001. Data are representative of two independent experiments.

### Molecular signature of *Giardia*-induced IL-10-producing Th2 cells at single cell resolution

After demonstrating that CD4^+^ T cells were the major source of *Giardia*-driven IL-10 in the small intestine of infected mice, our next step was to characterize the transcriptional program of these cells to understand their molecular nature at single cell resolution. For that, we used IL-10 GFP reporter mice and specifically sorted out IL-10^+^TCRβ^+^CD4^+^ T cells isolated from the lamina propria from both naïve and *Giardia*-infected mice (Fig.3A-B) to perform the single cell RNA sequencing analysis. The UMAP plot indicated the contribution of each cluster to the proportion of IL-10-producing CD4^+^ T cells from naïve and *Giardia*-infected mice.

**Figure 3:**
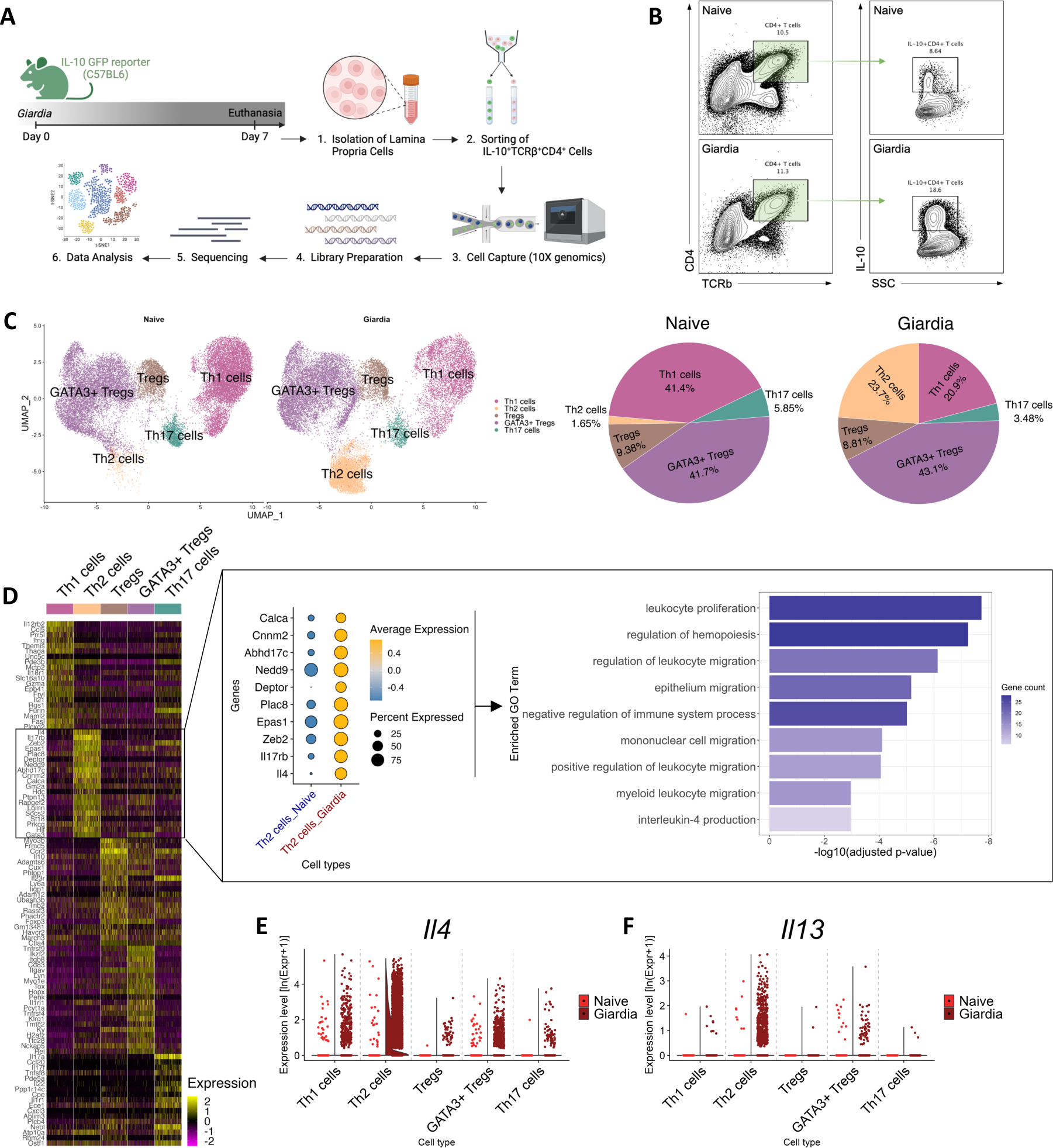
Single-cell transcriptional profiling of *Giardia*-induced IL-10-producing CD4^+^ T cells. (A) Experimental schematic. Small intestine lamina propria cells were isolated from *Giardia*-infected IL-10 GFP reporter mice (7 d.p.i.), and IL-10-producing CD4^+^ T cells were sorted for further analysis by single-cell RNA-seq. (B) Representative dot plots of sorted TCR3^+^CD4^+^IL-10^+^ from naïve and *Giardia*-infected mice (7 d.p.i.). (C) UMAP plots showing the identity of IL-10 producing CD4^+^ T cell subpopulations by RunUMAP and clustering analysis, and PoW plots indicating the frequency of T helper and T regulatory cell subpopulations in the naïve or *Giardia*-infected TCR3^+^CD4^+^IL-10^+^ sorted population. (D) Heatmap showing the expression of the most significantly 20 enriched transcripts in each subpopulation cluster and pathway enrichment analysis using the top 25 markers in Th2 cells. (E) and (F) Violin plots indicating IL-4 and IL-13 transcripts expression in CD4^+^ T cell subpopulations. Data are representative of one experiment using 8 naïve and 12 *Giardia*-infected mice.

Subpopulations of CD4^+^ T cells were annotated based on the gene expression level of the top10 highly upregulated signature genes (Supp.Fig.4A-B). Among the naïve samples, the most abundant populations of IL-10-producing CD4^+^ T cells consisted of GATA3^+^ Treg cells and Th1 cells (41.7% and 41.4%, respectively). Less abundant IL-10 producing populations included Tregs (9.38%), Th17 cells (5.85%), and lastly Th2 cells, representing only 1.65% of all IL-10-producing CD4^+^ T cells (Fig.3C). Analysis of the *Giardia*-infected mice demonstrated a marked expansion in the proportion of IL-10-producing Th2 effector cells to 23.7% (14-fold increase), with a significant contraction in the IL-10^+^ Th1 cluster to 20.9%. The proportion of the other subpopulations of CD4^+^ T cells remained similar to the naïve mice, with a contribution from the cluster of GATA3^+^ Treg cells around 43.1% (Fig.3C). This observation highlights the pivotal role these effector Th2 cells play in orchestrating the regulatory environment within the lamina propria during *Giardia* infection.

We further sought to unravel the transcriptional profiles of these expanding IL-10-producing Th2 cells. The gene expression analysis of the *Giardi*a-driven IL-10-producing Th2 cells demonstrated a significant upregulation of *Gata3*, *IL-17rb* (IL-25 receptor), *IL-4*, *Plac8*, *Calca*, *Epas1, IL-13*, and other genes when compared to IL-10^+^ Th2 cells from naïve mice (Fig.3D-F and Supp.Fig.4B and C). This transcriptional signature is commonly found in the pathogenic Th2 effector cells in models for allergic diseases and helminth infections (Gazzinelli-Guimaraes et al., 2023), suggesting that *Giardia* is inducing the expansion of polyfunctional Th2 effector cells that are not only a relevant source of Type 2 cytokines, but also an important source of IL-10 in the small intestine lamina propria (Supp.Fig.4C).

Through gene set enrichment analysis focusing on the top 25 genes expressed by this Th2 cell subset, we uncovered a multifaceted genetic profile closely linked to critical biological processes (Fig.3D). Notably, these processes encompassed the regulation of hemopoiesis and leukocyte migration, pivotal for the mobilization of immune effectors to the site of infection. Additionally, our analysis unveiled a significant association with epithelium migration, indicative of the dynamic cellular shifts essential for the immune surveillance and integrity of the intestinal barrier during infection. Intriguingly, the identified genetic profile was strongly associated with the negative regulation of immune system processes, highlighting the delicate equilibrium between immune activation and restraint that characterizes the host response to *Giardia* infection (Fig.3D). In sum, our findings intricately connect the dots between *Giardia* infection, IL-10-producing Th2 effector cells, and the underlying molecular signatures that define their functional role within the intestinal mucosa.

### Blockade of Type 2 immunity impairs *Giardia*-induced IL-10 and shifts mucosal immune response to a Type 1/17 phenotype that is associated with parasite clearance

To investigate the role of the *Giardia*-induced Th2 response in the orchestration of intestinal mucosal immunity and its implication on *Giardia intestinalis* parasitism, we infected STAT6 deficient (STAT6^-/-^) mice to compare their response against WT-infected mice (Fig.4A). The absence of STAT6 signaling during *Giardia* infection resulted in a significant reduction of *Giardia*-induced Th2 effector cells and IL-10^+^ Th2 cells in the small intestine lamina propria of infected mice, however it did not alter the number of IL-10 producing GATA3^+^ Tregs, suggesting that STAT6 signaling does not regulate the differentiation of GATA3^+^ Tregs in the intestine (Fig.4B). Further, *Giardia*-infected STAT6^-/-^ mice showed a proinflammatory shift marked by increased numbers of IL-17A^+^RORγt^+^ Th17 and IFNγ^+^Tbet^+^ Th1 effector cells (Fig.4C). This shift in the T helper cell polarization was also associated with a significant reduction of IL-4 and increase of IL-17A levels in the SI tissue homogenate. Strikingly, we observed a significant reduction in the levels of IL-10 in the small intestine of *Giardia*-infected mice deficient in STAT6 signaling when compared to WT (Fig.4D).

**Figure 4:**
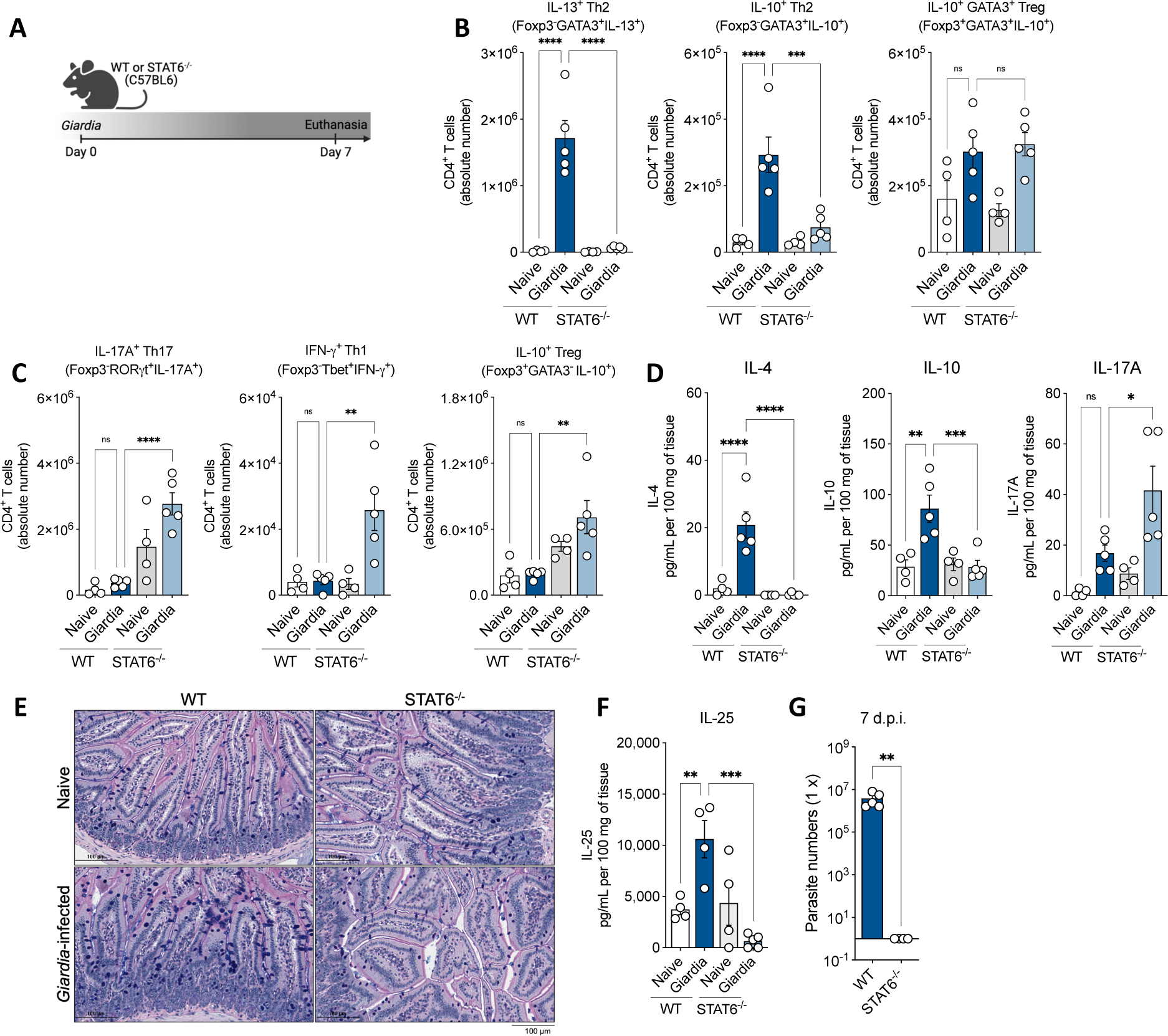
Type 2 signaling blockage impairs *Giardia*-induced IL-10 and shifts the mucosal immune response to a Type 1/17 phenotype that promotes parasite clearance. (A) Experimental schematic. Female WT or STAT6^-/-^ mice (C57BL/6 background) were perorally infected with 1×10^6^ *Giardia* GS/M strain trophozoites. (B) Scatter plot graphs indicating the absolute number of Th2 (Foxp3^-^GATA3^+^IL-13^+^), IL-10^+^ Th2 (FoxP3^-^GATA3^+^IL-10^+^), and IL-10^+^ GATA3^+^ Treg (FoxP3^+^GATA3^+^IL-10^+^) cells in the small intestine lamina propria of *Giardia*-infected mice (7 d.p.i.) (gated on Live CD45^+^TCR3^+^CD4^+^). (C) Scatter plot graphs indicating the absolute number of Th17 (FoxP3^-^RORγt^+^IL-17A^+^), Th1 (Foxp3^-^GATA3^-^Tbet^+^IFN-γ^+^), and IL-10^+^ Treg (FoxP3^+^GATA3^-^IL-10^+^) cells in the small intestine lamina propria of *Giardia*-infected mice (7 d.p.i.) (gated on Live CD45^+^TCR3^+^CD4^+^). (D) IL-4, IL-10, and IL-17A levels in the small intestine of WT or STAT6^-/-^ *Giardia*-infected mice (7 d.p.i.). (E) Representative image of the jejunum from WT or STAT6^-/-^ *Giardia*-infected mice stained with Alcian Blue/Periodic Acid Schiff (AB/PAS) for mucus production by goblet cells (7 d.p.i.). Scale bars represent 100 μm. (F) IL-25 levels in the small intestine tissue homogenate of WT or STAT6^-/-^ *Giardia*-infected mice (7 d.p.i.) measured by ELISA. (G) *Giardia* burden in the small intestine (duodenum) of WT or STAT6^-/-^ *Giardia*-infected mice (7 d.p.i.). Data are represented as mean ± SEM and significance was calculated with one-way ANOVA test followed by Sidak’s multiple comparisons test. *p≤0.05, **p≤0.01, ***p≤0.001. Data are representative of two independent experiments.

To further explore the role of Type 2 immunity induced by STAT6 signaling in the pathophysiology of the small intestine during *Giardia* infection, we assessed inflammation, mucus production, goblet cell hyperplasia, parasite burden, and IL-25 levels of WT versus KO mice. Our results demonstrated that the downstream impairment of the Th2 effector response, induced by the absence of STAT6 signaling in *Giardia*-infected mice, led to an increase in the infiltration of inflammatory cells to the tissue, along with a reduction of mucus production by goblet cells (Fig.4E), and a significant decrease in the production of IL-25 driven by *Giardia* (Fig.4F).

This reshaped mucosal response to a dominant Th17 inflammatory environment yielded compelling insights into the intricate dynamics between the regulatory role of Th2 effector responses, IL-10 induction, and *Giardia* parasitism within the small intestine. STAT6 deficient mice had dramatically reduced numbers of *Giardia* trophozoites infecting the small intestine, which led to a complete clearance of the infection by day seven (Fig.4G). This phenomenon was not observed in WT-infected mice, which continued to exhibit elevated numbers of parasites colonizing the gastrointestinal tract at the time point investigated (Fig.4G).

### Previous infection with *Giardia* induces protection against inflammatory bowel-like diseases

Inspired by the epidemiological observation that *Giardia* co-infection is associated with reduced inflammatory outcomes in bystander enteric infections (Bhavnani *et al*., 2012; Cotton *et al*., 2014b; Muhsen *et al*., 2014; Oberhelman *et al*., 2001; Veenemans *et al*., 2011; Wang *et al*., 2013), we next investigated the ability of *Giardia*-driven IL-10/Th2 axis in the immunoregulation of other intestinal inflammatory disorders. For this, we used two different IBD-like models. First, we developed a co-infection model by previously infecting WT mice orally with 1 x 10^6^ *Giardia intestinalis* trophozoites followed by oral infection with 10 cysts of *Toxoplasma gondii* three days later (Fig.5A). *T. gondii* is an intestinal protozoan parasite that drives an acute Th1/IFN-gamma-mediated lethal ileitis in C57Bl6/J mice that is a model for Crohn’s disease-like enteritis (Liesenfeld, 2002; Liesenfeld et al., 1996). Six to seven days post-*Toxoplasma* infection, we observed that mice singly infected with *T. gondii* or co-infected with *Giardia* lost about 10-12% of their body weight, whereas naïve and *Giardia* singly infected mice did not (Fig.5B). Strikingly, histopathological analysis demonstrated that *Toxoplasma*-infected mice that had been previously infected with *Giardia* presented with a remarkably reduced inflammatory infiltrate in the small intestine when compared to single *Toxoplasma*-infected mice (Fig.5C). Additionally, prior infection with *Giardia* downregulated the levels of IFN-γ in the small intestine of *Toxoplasma* co-infected mice, while significantly increasing IL-10 levels (Fig.5D). Along with these results, the presence of *Giardia* reduced the frequency of Tbet^+^IFNγ^+^ Th1 cells in the SI LP and prevented the Treg (FoxP3^+^) and GATA3^+^ Treg collapse in *Toxoplasma* co-infected mice eight days-post *Toxoplasma* infection (8 d.p.T.i.) (Fig.5E) – Th1 and Treg are the major hallmarks of the *Toxoplasma*-driven pathogenesis in the small intestine (Oldenhove et al., 2009). Interestingly, the downregulation of the Type 1 response and the increased IL-10-driven regulatory response elicited by *Giardia* parasites allowed for greater *T. gondii* proliferation and a significantly increased parasite burden without associated sequelae at seven days post *T. gondii* infection (Supp.Fig.5A).

**Figure 5:**
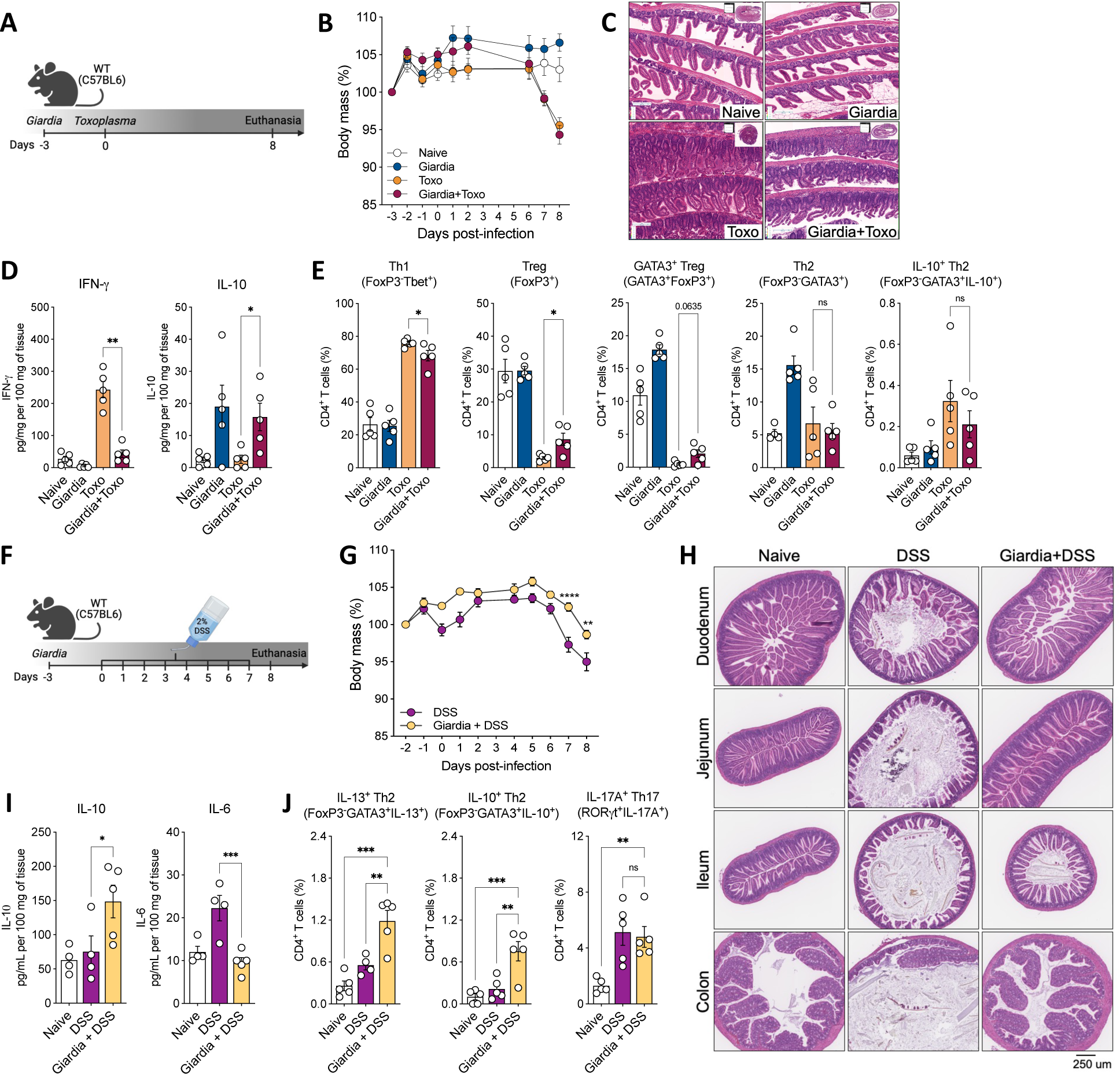
Previous infection with *Giardia* induces protection against inflammatory bowel-like diseases. (A) Experimental schematic. Female C57BL/6 mice were perorally infected with 1×10^6^ *Giardia* GS/M strain trophozoites and three days later mice were perorally co-infected with 10 cysts of *Toxoplasma gondii* Type II 76K GFP-Luc strain. (B) Body weight loss of mice single infected with *Toxoplasma* or co-infected with *Giardia* and *Toxoplasma* were monitored daily. (C) Representative image of hematoxylin-and-eosin staining (H&E) of the proximal small intestine (swiss roll) from naïve, *Giardia*-, *Toxoplasma*– or *Giardia+Toxoplasma*-infected mice (8 days-post *Toxoplasma* infection, 8 d.p.T.i.). Scale bars represent 160 μm. (D) IFN-γ and IL-10 levels in the small intestine of singly or co-infected mice (8 d.p.T.i.) measured by Luminex. (E) Scatter plot graphs indicating the frequency of Th1 (FoxP3^-^Tbet^+^), total Treg (FoxP3^+^), GATA3^+^ Treg (GATA3^+^FoxP3^+^), Th2 (GATA3^-^Foxp3^+^), and IL-10^+^ Th2 (FoxP3^-^GATA3^+^IL-10^+^) cells in the small intestine lamina propria of singly or co-infected mice (8 d.p.T.i.). Gated on Live CD45^+^TCR3^+^CD4^+^. (F) Experimental schematic. Female C57BL/6 mice were perorally infected with 1×10^6^ *Giardia* GS/M strain trophozoites and three days later mice were administered 2% DSS drinking water or normal drinking water for 7 consecutive days. From day 7 to day 8 mice were placed back on normal drinking water, and euthanasia was performed on day 8 (8 days-post treatment, 8 d.p.t.). (G) Body weight loss of DSS-treated mice infected or not with *Giardia* was monitored daily. (H) Representative image of H&E staining of the small and large intestine sections from DSS-treated mice infected or not with *Giardia*. Scale bars represent 250 μm. (I) IL-10 and IL-6 levels in the distal small and proximal large intestines of DSS-treated mice infected with or without *Giardia* (8 d.p.t.) measured by Luminex. (J) Scatter plot graphs indicating the frequency of Th2 (FoxP3^-^GATA3^+^IL-13^+^), IL-10^+^ Th2 (Foxp3^-^GATA3^+^IL-10^+^), and Th17 (RORγt^+^IL-17A^+^) cells in the colonic lamina propria of DSS-treated mice infected or not with *Giardia* (8 d.p.t.). Gated on Live CD45^+^TCR3^+^CD4^+^. Data are represented as mean ± SEM and significance was calculated with one-way ANOVA test followed by Sidak’s multiple comparisons test. *p≤0.05, **p≤0.01, ***p≤0.001. Data are representative of three (A-E) and two (G-J) independent experiments.

Importantly, at this later point of *Toxoplasma* infection (8 d.p.T.i.), we did not observe differences in the Th2 compartment, verified by the equal frequency of Th2 (FoxP3^-^GATA3^+^) and IL-10-producing Th2 (FoxP3^-^GATA3^+^IL-10^+^) cells driven by *Giardia* in the co-infected group when compared to the *Toxoplasma* singly infected mice (Fig.5E). However, in an earlier time point (4 d.p.T.i.) in the co-infection model, before the establishment of the strong and dominant *Toxoplasma*-driven Th1 response, we did observe that *Giardia* was capable of inducing a high frequency of Th2 and IL-10-producing Th2 cells in the SI LP of co-infected mice when compared to *Toxoplasma* singly infected group (Supp.Fig.5B).

We next evaluated the capacity of a prolonged *Giardia* infection (4 weeks post-infection) to elicit a regulatory immune response (specifically, IL-10 production) and to ameliorate intestinal inflammation induced by *Toxoplasma gondii*. Notably, our findings revealed that mice with prolonged *Giardia* infection, subsequently infected with *Toxoplasma*, exhibited sustained, elevated levels of IL-10. This increase was linked to a significant reduction in the inflammatory score of the intestine, as compared to mice infected solely with *Toxoplasma*. Remarkably, the effect observed four weeks post-*Giardia* mirrored the response noted seven days post-infection (Supp.Fig.5C and D).

Since *Giardia* also drives a Th2 immune response in the colon, we developed a Th1/Th17 acute colitis model driven by the administration of DSS for seven consecutive days in *Giardia*–previously infected mice (three days before the DSS treatment started) (Fig.5F). Interestingly, the presence of *Giardia* reduced the percentage of weight loss in DSS-induced colitis mice when compared to those that developed colitis but were not infected (Fig.5G). Remarkably, the histological analysis demonstrated that previous infection with this gastrointestinal protozoan parasite ameliorated DSS-driven inflammation and attenuated the colitis outcome in all sections of the small intestine and colon of mice compared against the non-infected DSS-induced colitis group (Fig.5H). In addition, the presence of *Giardia* in mice that developed acute colitis was associated with enhanced IL-10 and decreased IL-6 levels in the distal small intestine (Fig.5I). Finally, although *Giardia* infection did not alter the number of IL-17A^+^ Th17 cells in the colon of mice with colitis, previous infection with the parasite significantly increased the number of polyfunctional IL-13 and IL-10 producing Th2 effector cells in mice with DSS-induced acute colitis (Fig.5J). Taken together, these data substantiate the epidemiological observations, and demonstrate that the IL-10/Th2 axis driven by *Giardia* in the gastrointestinal tract plays a role in regulating bystander intestinal inflammation.

### *Giardia*-mediated IBD protection is dependent on STAT6 signaling-Th2 response axis

Because *Giardia* infection attenuated bystander intestinal inflammatory responses by 1) suppressing the induction of IFN-γ producing Tbet^+^ Th1 cells during co-infection with *T. gondii,* and 2) reducing IL-6 in DSS-induced colitis model, we next investigated the mechanisms associated with such protection. We interrogated whether *Giardia*-driven Type 2 mucosal immunity plays a role in the observed protection during inflammatory bowel-like diseases. For that, we repeated the IBD models, but now using the STAT6 deficient mice in the context of *Giardia* infection (Fig.6A and F). For the *Toxoplasma*-driven acute ileitis model, although both co-infected groups showed a relevant weight loss from six to seven days-post *T. gondii* infection, the weight loss was significantly greater in the STAT6 deficient mice compared to WT (Fig.6B). This observation was associated with increased histological inflammation in the small intestine in the STAT6^-/-^ co-infected mice (Fig.6C). Furthermore, we observed exacerbated *T. gondii*-mediated IFN-γ and IL-6 production, associated with decreased IL-13, in the SI tissue when the STAT6 signaling was absent during *Giardia* co-infection, while IL-10 levels were not significantly (p=0.0519) altered at this time point in both co-infected groups (Fig.6D). Notably, linked with this more prominent proinflammatory response, we also observed an increased frequency of FoxP3^-^Tbet^+^ Th1 cells and a significant decrease in Treg (FoxP3^+^), GATA3^+^ Treg (FoxP3^+^GATA3^+^), Th2 (FoxP3^-^GATA3^+^), and IL-10-producing Th2 (FoxP3^-^GATA3^+^IL-10^+^) cells in the co-infected STAT6^-/-^ mice when compared to co-infected WT mice (Fig.6E), suggesting that the absence of STAT6 signaling failed to suppress the *Toxoplasma* driven type-1 inflammatory response and its downstream effect on tissue inflammation during *Giardia* infection.

**Figure 6:**
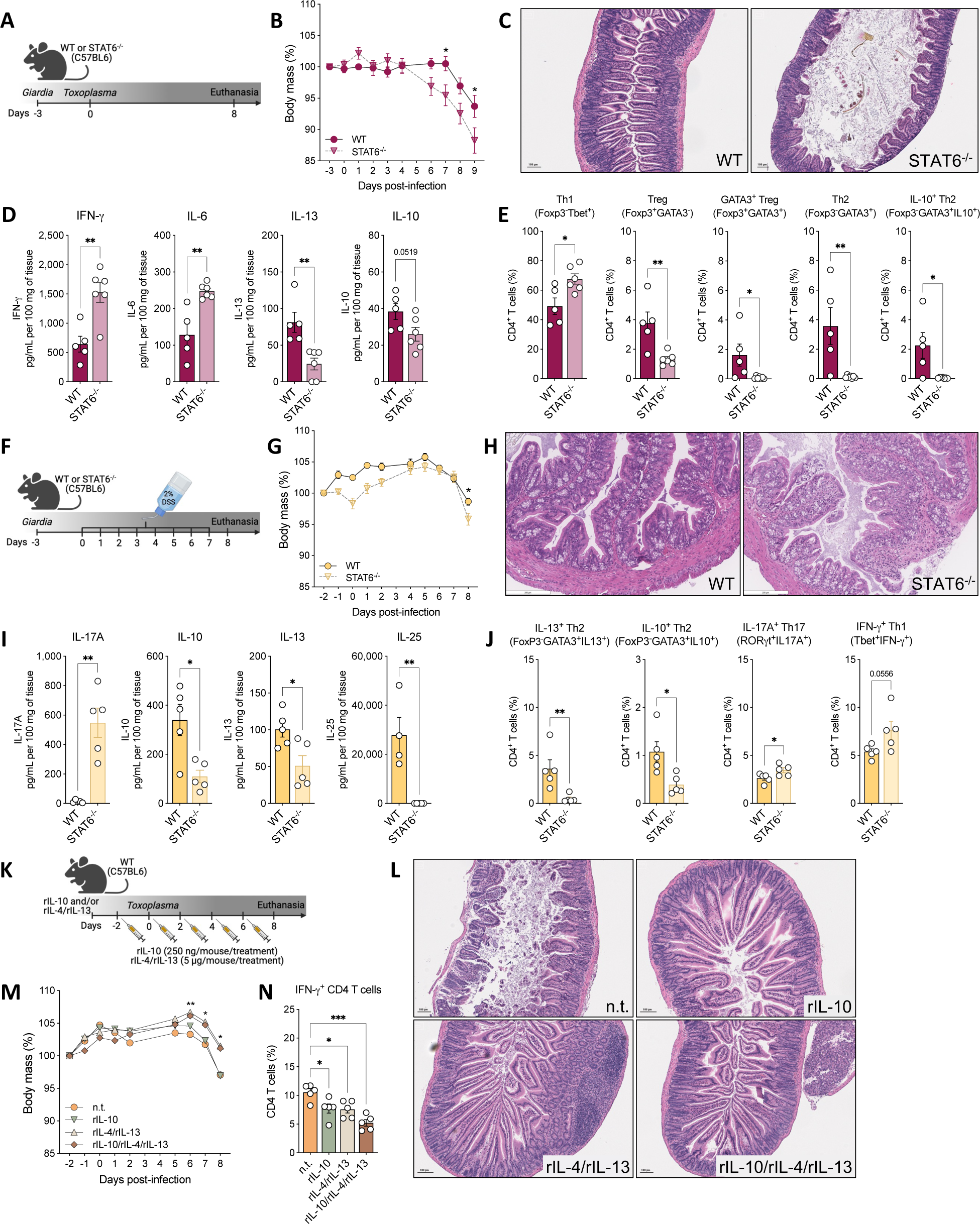
Type 2 signaling blockage reduces *Giardia*-mediated protection against inflammatory bowel-like diseases. (A) Experimental schematic. Female WT or STAT6^-/-^ mice (C57BL/6 background) were perorally infected with 1×10^6^ *Giardia* GS/M strain trophozoites and three days later mice were perorally co-infected with 10 cysts of *Toxoplasma gondii*. Euthanasia was performed 8 days post-*Toxoplasma* infection (8 d.p.T.i.). (B) Body weight loss of WT or STAT6^-/-^ mice co-infected with *Giardia* and *Toxoplasma* were monitored daily. (C) Representative image of H&E staining of the proximal small intestine from WT or STAT6^-/-^ mice co-infected with *Giardia* and *Toxoplasma* (8 d.p.T.i.). (D) IFN-γ, IL-6, IL-13, and IL-10 levels in the proximal small intestine of WT or STAT6^-/-^ co-infected mice (8 d.p.T.i.) measured by Luminex. (E) Scatter plot graphs indicating the frequency of Th1 (Foxp3^-^Tbet^+^), Treg (Foxp3^+^GATA3^-^), GATA3^+^ Treg (GATA3^+^FoxP3^+^), Th2 (GATA3^-^Foxp3^+^), and IL-10^+^ Th2 (FoxP3^-^GATA3^+^IL-10^+^) cells in the small intestine lamina propria of WT or STAT6^-/-^ co-infected mice (8 d.p.T.i.). Gated on Live CD45^+^TCR3^+^CD4^+^. (F) Experimental schematic. WT or STAT6^-/-^ mice were perorally infected with 1×10^6^ *Giardia* GS/M strain trophozoites and three days later mice were administered 2% DSS drinking water or normal drinking water for 7 consecutive days. From day 7 to day 8 mice were placed back on normal drinking water, and euthanasia was performed on day 8 (8 d.p.t.). (G) Body weight loss of WT or STAT6^-/-^ *Giardia*-infected and DSS-treated mice was monitored daily. (H) Representative image of H&E straining of the colon from *Giardia*-infected and DSS-treated mice. Scale bars represent 200 μm (8 d.p.t.). (I) IL-17, IL-10, IL-13, and IL-25 levels in the distal small and proximal large intestines of WT or STAT6^-/-^ *Giardia*-infected mice and treated with DSS (8 d.p.t.) measured by Luminex. (J) Scatter plot graphs indicating the frequency of Th2 (FoxP3^-^GATA3^+^IL-13^+^), IL-10^+^ Th2 (Foxp3^-^GATA3^+^IL-10^+^), Th17 (RORγt^+^IL-17A^+^), and Th1 (Tbet^+^IFN-γ^+^) cells in the colonic lamina propria of WT or STAT6^-/-^ *Giardia*-infected mice and treated with DSS (8 d.p.t.). Gated on Live CD45^+^TCR3^+^CD4^+^. (K) Experimental schematic. Female C57BL/6 mice were perorally infected with 1×10^6^ *Giardia* GS/M strain trophozoites (day –3) or treated with recombinant IL-10 (250 ng/mouse) and/or recombinant IL-4 and IL-13 (5ug/mouse each) every two days (days 0, 2, 4, and 6), and on day 0 all groups were perorally infected with 10 cysts of *Toxoplasma gondii*. Euthanasia was performed 8 days post-*Toxoplasma* infection (8 d.p.T.i.). (L) Representative image of H&E staining of the proximal small intestine from *Toxoplasma*-infected mice that were co-infected with *Giardia* or treated with rIL-10 and/or rIL-4/rIL-13 (8 d.p.T.i.). (M) Body weight loss of mice was monitored daily. (N) Scatter plot graph indicating the frequency of IFN-γ^+^ CD4^+^ T cells in the small intestine lamina propria of *Toxoplasma*-infected mice that were co-infected with *Giardia* or treated with rIL-10 and/or rIL-4/rIL-13 (8 d.p.T.i.). Gated on Live CD45^+^TCR3^+^CD4^+^. Data are represented as mean ± SEM and significance was calculated with non-parametric Mann-Whitney test (D-E, I-J) and one-way ANOVA test followed by Sidak’s multiple comparisons test (B, M, N). *p≤0.05, **p≤0.01, ***p≤0.001. Data are representative of two independent experiments.

We also used the DSS-induced acute colitis model in *Giardia*-infected WT and STAT6^-/-^ mice (Fig.6F), and although both groups showed reduction in the body weight five to six days-post DSS administration, we observed that STAT6 deficient mice had greater weight loss at day 8 compared to WT (Fig.6G). Histopathological analysis indicated that the absence of STAT6 signaling was associated with increased inflammatory infiltrate in *Giardia*-infected mice that developed acute colitis, while this was not observed in the WT group (Fig.6H). Additionally, we observed increased IL-17A and decreased IL-10, IL-13, and IL-25 levels in the distal small intestine (Fig.6I), alongside a reduction in the frequency of IL-13^+^ Th2 effector cells and IL-10^+^ Th2 cells in *Giardia*-infected STAT6^-/-^ mice with colitis (Fig.6J). Lastly, there was an increased frequency in the IL-17A^+^ Th17 and IFN-γ^+^ Th1 cells in the absence of STAT6 signaling (Fig.6J).

Finally, to test whether the IL-10/Th2 response induced during *Giardia* infection is sufficient to confer protection against bystander intestinal inflammation, we treated *Toxoplasma*-infected WT mice with either recombinant IL-10 and/or IL-4/IL-13 cytokines every other day for 10 days (Fig.6K). The *Toxoplasma*-infected non-treated (n.t.) and rIL-10 treated groups showed greater weight loss compared to the mice treated with rIL-4/IL-13 or rIL-10/rIL-4/rIL-13 (Fig.6M). The histopathological analysis showed reduced inflammation and less tissue damage in the proximal small intestine of all treated mouse groups (Fig.6L), along with decreased frequency of IFN-γ^+^ CD4^+^ T cells in the SI LP (Fig.6N). The protection following treatment with recombinant IL-4, IL-10, and IL-13 appeared to be partial, suggesting that the presence *Giardia* and its induced response may be essential for comprehensive protection.

Taken together, our data indicate that the *Giardia*-triggered Th2 immune response in the gastrointestinal tract, mediated by STAT6 signaling and Type 2 cytokines, is instrumental in upholding the intestinal integrity and homeostasis. This response culminated in increased IL-10 levels within the small intestine, serving a crucial role in shielding against the immunological challenges posed by Type 1– and Type 17-induced inflammatory bowel diseases.

## DISCUSSION

In this study a significant reshaping of intestinal mucosal immunity occurred during infection with the gastrointestinal protozoan parasite *Giardia intestinalis*. This parasite is highly prevalent worldwide and was identified as a common enteric parasite in our Nigerian cohort of school-aged children. In our mouse model of infection, an initial Th17 response, commonly induced by extracellular pathogens, was replaced by a significant and substantial Type 2-associated cytokine response (IL-4 and IL-13) which promoted chronic carriage and an increase in cyst load, the transmissible stage of the parasite. This shift toward a dominant Type 2 immune response was accompanied by various changes within the host, including elevated levels of antigen-specific Th2 effector cells, an enhanced production of IL-10, heightened goblet cell activity and increased mucus secretion, as well as elevated IL-25 levels in the small intestine. When Th2 signaling was disrupted using STAT6^-/-^ mice, a noticeable shift away from Type 2 induced immunity led to the emergence of Th1 and Th17 phenotypes that controlled *Giardia* proliferation, but was associated with collateral damage, specifically the induction of a dysregulated pathophysiology in the small intestine, as observed previously (Singer and Nash, 2000). Our results suggest that any regulation in the development of the dominant Th2 immune response negatively impacts parasite persistence, transmissibility and promotes disease pathogenesis.

Investigation into the immune landscape post-*Giardia* infection, at single cell resolution, using both a multiparameter immunophenotyping strategy along with functional characterization and molecular profiling by single cell RNA sequencing, highlighted the significance of the *Giardia*-driven IL-10 producing FoxP3^-^GATA3^+^ Th2 effector cells in regulating the immune environment within the host. The transcriptional signature of this IL-10^+^ Th2 cells was synonymous with a pathogenic subset of memory Th2 cells (termed Th2A or Tpath2 cells) that show upregulated expression of *Gata3*, *Il17rb*, *Plac8*, *Calca*, *Il4* and *Il13*. Indeed, the heterogeneity of these distinct Th2 effector cells subsets, including their capacity to produce large amounts of different cytokines and to differentiate into a vast repertoire of T effector cell types, has been the subject of extensive studies only recently. Interestingly, these subsets of polyfunctional Tpath2 effector cells that produce large amounts of IL-10, as observed during *Giardia* infection, are common to inflammatory responses driven by allergens and helminth parasites (Gazzinelli-Guimaraes et al., 2023), which are known to impact the pathogenesis of Type 2 inflammatory responses. Indeed, Metenou and collaborators (Metenou et al., 2010) showed that in chronically infected individuals with filarial parasites, the prevalence of IL-10-mediated regulation in helminth infection primarily relies on the activity of IL-10 producing Th2 effector cells.

Understanding the regulatory effects of Th2 responses, and specifically how reduced Th2 signaling significantly affects IL-10 induction and the expansion of immune cells, has the potential for managing intestinal parasitic infections more effectively and holds the promise for therapeutic intervention. *Ex vivo* studies have shown that excreted/secreted (ES) antigens of *Giardia muris* (the murine *Giardia* species) can induce high levels of IL-4 and IL-5 from Peyer’s patch (PP) isolated cells from infected mice during *Giardia* infection (Djamiatun and Faubert, 1998; Petro et al., 1992). These studies also established that antigenic stimulation of splenocytes from *Giardia intestinalis*-infected BALB/c mice induced elevated levels of IL-4, IL-5 and IL-10 (Jimenez et al., 2014; Jimenez et al., 2009). Stimulation of splenocytes and mesenteric lymph node cells from *Giardia intestinalis*-infected C57BL/6 mice also showed elevated levels of IL-13, IL-17, IL-10 and IL-22 (Solaymani-Mohammadi and Singer, 2011), and RT-PCR studies identified elevated levels of IL-17A in the small intestine itself (Dann *et al*., 2015; Dreesen *et al*., 2014). Furthermore, it has previously been shown that PBMCs from patients that were exposed to *Giardia* infection during their lifetime upregulate the production of IL-10 and IL-13 levels after *in vitro* stimulation with *Giardia* protein lysate, when compared with stimulated PBMCs from healthy donors (Saghaug et al., 2016). This body of work provides a rationale to support the epidemiological observations that *Giardia* infection, by its ability to promote Type 2 immunity, is associated with reduced risk of severe diarrheal disease in children during infection with proinflammatory viruses and protists (Bhavnani *et al*., 2012; Cotton *et al*., 2014b; Muhsen *et al*., 2014; Oberhelman *et al*., 2001; Veenemans *et al*., 2011; Wang *et al*., 2013).

The importance of Type 2 immune responses in the pathophysiology of helminth infection has been instrumental in demonstrating how the administration of helminth excretory-secretory (ES) products or helminth infection can effectively mitigate inflammation in a variety of inflammatory disease models, including the DSS-induced colitis model in mice, as highlighted in the comprehensive review by Gazzinelli-Guimaraes and Nutman (Gazzinelli-Guimaraes and Nutman, 2018). Specifically, it has been established that ES products from *Ancylostoma ceylanicum* (a hookworm affecting humans, cats, dogs, and rodents) (Cancado et al., 2011), *Ancylostoma caninum* (a canine hookworm) (Ferreira et al., 2013), and *Trichinella spiralis* (a roundworm in carnivorous animals) (Yang et al., 2014) all alleviate the severity of DSS-induced colitis in mouse models. Additionally, extracellular vesicles from *Nippostrongylus brasiliensis* and *Trichuris muris* have exhibited modulatory effects on colitis in experimental models (Eichenberger et al., 2018). A common mechanism across these studies is the observation that elevated levels of Type 2-associated and regulatory cytokines, such as IL-10 and TGF-β, are associated with a marked decrease in proinflammatory cytokines including IL-6, IL-1β, IFN-γ, and IL-17A. These proinflammatory cytokines are known to contribute to the pathology seen in colitis and impact worm fecundity.

We evaluated the ability of *Giardia* infection to downregulate bystander intestinal inflammatory responses using two different IBD models, a Th1/Th17 colitis driven by dextran sodium sulfate (DSS) and a Th1-driven/IFN-gamma-mediated lethal ileitis induced by *Toxoplasma gondii*, an intestinal protozoan parasite that causes a Crohn’s disease-like enteritis. In both models we observed reduced inflammation in the intestines of *Giardia*-infected mice and this was associated with regulation of proinflammatory markers, including the suppression of IFN-γ producing Tbet^+^ Th1 cells during co-infection with *T. gondii*; and reduced IL-6 and increased IL-10 and IL-13 levels in DSS-induced colitis. More importantly, our results suggest that these protective mechanisms were dependent on *Giardia*-driven Th2 immunity and STAT6 signaling in the intestine, as *Giardia*-infected STAT6^-/-^ mice no longer regulated collateral damage induced during the development of ileitis or colitis.

Mechanisms by which *Giardia* influences the differentiation of CD4^+^ T cell responses are not well defined. Kamda and Singer showed that parasite extracts block the production of IL-12 by bone marrow derived dendritic cells (DCs) activated with Toll-like receptors (TLR) agonists (Kamda and Singer, 2009). A similar study with human DCs showed that ammonia, a product of parasite secreted arginine deiminases also reduced the production of IL-12p40 by DCs activated with LPS (Banik et al., 2013). *In vivo*, *Giardia* infection was shown to reduce IL-12p40 levels in the colons of mice treated with Cdiff toxin (TcdAB) (Cotton *et al*., 2014b). Secretion of cysteine proteases by the parasite has been implicated in reduced levels of IL-8 and reduced neutrophil recruitment (Cotton et al., 2014a). Specifically how *Giardia* promotes mucosal immunity reprogramming to facilitate both its life cycle progression and limit the collateral damage from Th1/Th17 inducing agents has not been defined. In studies of other protists and helminths, parasite products are known to target specific host pathways to modulate host immunity and facilitate parasitism. For example, *Tritrichomonas musculus* induces Type 2 immunity in the small intestine lamina propria of mice by secreting a protist metabolite, succinate, that triggers Tuft cell activation via SUCNR1 receptor, culminating in IL-25 secretion and ILC2 activation, which produces IL-13 and initiates the Type 2 immune response in the gut (Howitt et al., 2016; Lei et al., 2018; Nadjsombati et al., 2018; Schneider et al., 2018). It was also demonstrated that colonization with *Tritrichomonas* in the mouse cecum confers protection against lethal challenges of mucosal bacterial infections, such as *Salmonella typhimurium,* but the heightened inflammation during commensal carriage increases the risk of colitis and colorectal cancer (Chudnovskiy et al., 2016; Escalante et al., 2016). In addition, it has been shown that the intestinal murine nematode *Heligmosomoides polygyrus* express TGM1, a parasite factor that structurally mimics the TGF-β cytokine. TGM1 binds to the host receptor for TGF-β in naïve T lymphocytes to induce FoxP3 expression and promotes the differentiation of immunosuppressive Tregs as a mechanism to suppress intestinal inflammation and evade the host immunity during infection (Johnston et al., 2017). Additionally, it was demonstrated that the administration of TGM1 suppress DSS-induced colitis by reducing inflammation in mice, although it did not fully block the development of intestinal pathology (Smyth et al., 2021; Smyth et al., 2023). Lastly, another parasite factor shown to modulate host immunity is the *Schistosoma mansoni* egg-secreted glycoprotein T2 ribonuclease/omega-1. Omega-1 acts directly on DCs and suppresses TLR-dependent activation, altering the ability of DCs to form antigen-dependent conjugates with CD4^+^ T cells. This results in weak TCR signaling that mimics the effect of a low-dose antigen, which promotes Th2 polarization, and the formation of granulomas, promoting chronic carriage of paired schistosome worms and egg production (Steinfelder et al., 2009).

This study raises and supports the hypothesis that *Giardia*-derived molecules are actively promoting a Type 2 dominated environment to *Giardia*’s advantage, which facilitates parasitism and the transmissibility of the parasite’s life cycle. In so doing, it reshapes mucosal immunity to confer a mutualistic protection against bystander intestinal inflammation, protecting the host from Th1/Th17-driven pathologies induced by proinflammatory pathogens and chemical agents that induce colitis. In conclusion, the present study sheds new lights on the substantial impact of *Giardia* infection on reshaping mucosal immunity toward a regulatory Type 2 environment and provides insights into potential strategies for modulating mucosal immune responses to improve outcomes from wide variety of proinflammatory triggers that influence epithelial barrier integrity, dysbiosis, inflammation and disease development.

## MATERIAL AND METHODS

### Ethics statement

Ethical approval was obtained from the ethics committees of the State Ministries of Health and Hospital Management Boards of Nigeria for human stool samples human collection. A written consent was obtained from all participants. Protocols number: ESUTHP/C-MAC/RA/034/VOL 3/120 (ESUT Teaching Hospital, Enugu), CRS/MH/HREC/022/Vol. 1/216 (Ministry of Health, Cross River state), NHREC/09/23/2010B (Plateau State Specialist Hospital Jos), NHREC/17/03/2030 (Ministry of Health, Kano state), HMB/OFF/215/VOL II/564 (Benue State Hospital Management Board) and MOH/SEC3/1/60 (Ministry of Health Jigawa state).

### Sample Collection

Participants were given sample bottle labelled with unique ID to provide fresh stool samples. A questionnaire was administered to each participant to collect demographic and socioeconomic data. A total of 664 freshly collected stool samples from six different states in Nigeria were mixed individually with an equal volume of 95% ethanol within an hour to preserve the integrity of the DNA. Samples were shipped immediately to the Molecular Parasitology Section at the Laboratory of Parasitic Diseases, National Institute of Allergy and Infectious Diseases, NIH, USA for molecular analysis.

### DNA Isolation

DNA was isolated using the QIAsymphony SP equipment, which is intended for automated nucleic acid purification in molecular diagnostic applications. The Qiagen QIAsymphony PowerFecal Pro DNA Kit was used to pretreat the samples prior to loading on the QIAsymphony according to manufacturer’s instructions. After pretreatment, the supernatant was transferred to sterile 2 mL microtubes and placed on the QIAsymphony instrument for the bind DNA, washing and elution steps. The eluate was collected in a 96-well plate with the elution volume set at 100 μL.

### Real Time PCR Detection

A 62 bp fragment of the SSU RNA gene was amplified by real time PCR using forward (*Giardia*-80F – 5′-GACGGCTCAGGACAACGGTT-3′) and reverse (*Giardia*-127R – 5′-TTGCCAGCGGTGTCCG-3′) primers (Verweij et al., 2004). For each experiment, 2 μL of template DNA was added to qPCR assays comprising 5 μL of SYBR green and 0.1 pM of each primer in a final volume of 10 μL. Each experiment was performed in duplicates, and appropriate negative and positive controls were included. The amplification parameters were as follows, 10 min at 95 °C followed by 40 cycles of 10 sec at 95 °C and 40 sec at 60 °C using the Quant 6 real-time PCR instrument (Applied Biosystems). Samples were considered positive for *Giardia* spp. if the mean CT (threshold cycle) values were < 35.

### Mice

Wild-type (WT) C57BL/6 (strain 000664) and IL-10 GFP-reporter mice (strain 014530) on a C57BL/6 genetic background (B6(Cg)-*Il10^tm1.1Karp/J),^* females, 6-7 weeks-old, were purchased from The Jackson Laboratory (Bar Harbor, ME). STAT6 knockout (KO) mice on a C57BL/6 genetic background (females, 7 to 10 weeks-old) were kindly provided by Dr. P’ng Loke from the Laboratory of Parasitic Diseases, National Institute of Allergy and Infectious Diseases, National Institutes of Health. All mice were bred and maintained at the NIAID/NIH Animal Facility under specific pathogen free (SPF) conditions in accordance with the American Association for the Accreditation of Laboratory Animal Care (AAALAC). All experiments were performed under the Animal Study Protocol LPD-22E, approved by NIAID Animal Care and Use Committee (ACUC).

### *Giardia intestinalis* culture, *in vivo* infection and parasite burden

*Giardia intestinalis* GS/M (clone H7) strain trophozoites were obtained from ATCC (catalog no. 50581) and cultivated *in vitro* for *in vivo* infections as previously described by Fink and collaborators (Fink et al., 2020). Briefly, parasites were cultured in TYI-S-33 medium supplemented with 10% adult bovine serum, 10 U/mL Penicillin/Streptomycin, and 250 ng/mL Amphotericin B (Thermo Fisher Scientific). For experimental infections, *Giardia* trophozoites were harvested in cold PBS, washed twice, and 1 x 10^6^ *Giardia* trophozoites in 0.2 ml of PBS were used to infect mice via gavage. For the assessment of *Giardia* burden *in vivo*, a section of 3 cm of the duodenum (immediately distal to the ligament of Treitz, about 2 to 3cm below the stomach) was collected and incubated in 4 mL of PBS on ice for at least 30 min. Intestinal sections were gently scraped, and trophozoites were counted on a hemocytometer.

### Toxoplasma gondii maintenance and in vivo infection

Cysts from the type II *Toxoplasma gondii* strain 76K, expressing green fluorescent protein and firefly luciferase (76K-GFP/Luc), were maintained in wild-type C57BL/6 mice by oral passage of five cysts/mouse. For the experimental infections and co-infections, mice were infected with 10 cysts of 76K-GFP/Luc strain obtained from *T. gondii*-infected mouse brains homogenate. Eight days-post *T. gondii* infection, mice were euthanized to assess the immunological response.

### DSS-induced colitis

Acute colitis was induced in naïve or *Giardia*-infected C57BL/6 WT mice by 2% (w/v) Dextran Sulfate Sodium Salt (DSS colitis grade, M.W. 36,000-50,000, MP Biomedicals) added to the drinking water, as described by Wirtz and collaborators (Wirtz et al., 2017). Mice were orally infected with 1 x 10^6^ *Giardia* trophozoites, and three days later DSS administration begin. Mice were left on 2% DSS water for seven consecutive days (0 to 7), and then they were placed on normal drinking water for one day (7 to 8). On day eight, mice were euthanized, and the colitis severity was assessed by tissue histology, inflammatory cytokine levels, and flow cytometry.

### Isolation of Lamina Propria Cells

Lamina propria cells were isolated from small and large intestine sections as previously described (Kim et al., 2022). Briefly, the intestinal sections were excised, washed with cold Hank’s Balanced Salt Solution (HBSS) supplemented with 5% FBS and 25 mM HEPES (5% HBSS), followed by Peyer patches removal and longitudinally opening of the intestinal segments. Intestines were then cut into 2 cm fragments and washed once in 5% HBSS, followed by one wash with HBSS with 1.3 mM EDTA. Small intestine fragments were then incubated in a shaker (80 RPM) for 15 min at 37°C with 5% HBSS containing 1 mM DTT to remove luminal mucous. After the incubation with DTT, the intestinal fragments were washed five times with cold 5% HBSS, followed by tissue digestion for 1 h at 37°C with Iscov’s digestion media containing 1 mg/ml Liberase TL (Roche), and 0.5 mg/ml DNAse (Thermo) (supplemented with 10% FBS, 5% NCTC 109, 10 mM HEPES, 1% Glutamax, 10 U/mL Penicillin/Streptomycin, 0.1% β-mercaptoethanol) in the shaking incubator (80 RPM). Finally, tissues were passed through a 70 μm cell strainer, and the lamina propria cells were stimulated *in vitro* for the antigen-specific immune response assay or immunophenotypic analysis by flow cytometry.

### Flow cytometry

For the phenotypic and functional characterization of the small intestine at T cell level, small intestine lamina propria (SI LP) cells were incubated for 2.5 hours at 37°C, 5% CO_2_ upon stimulation with 0.5/0.05 nM of PMA/ionomycin (Sigma-Aldrich) and 10 μg/mL of brefeldin A. Briefly, staining using the T cell polarization panel required 3 steps: The cells were stained with a LIVE/DEAD marker (Fixable Blue, UV450, Thermo Fisher Scientific). Secondly, after washing with FACS buffer (Staining buffer, Biolegend, USA), antibodies against the extracellular markers CD45, TCR-β, CD90.2, CD19, CD11b, ST2, CD4 were used. After washing, cells were then fixed with 2% paraformaldehyde and permeabilized (Perm buffer, eBioscience), and finally stained with the following intracellular antibodies: GATA-3, T-bet, RORγt, FoxP3, IL-10, IL-13, IL-17A, and IFN-γ. A description of the antibodies used for the T cell panel, including the source and catalog/clone numbers, are included in Table 3. Data were acquired on an LSR Fortessa flow cytometer (BD Biosciences) and analyzed with FlowJo software (Tree Star) using the gating strategy 1 (Supp.Fig.3A-J). Then, to determine the source of IL-10 driven by *Giardia* infection, lamina propria cells were isolated from the SI LP of naïve and infected IL-10 GFP-reporter mice and were then stained for a myeloid and lymphoid lineage panel (see Table 3), using a 2-step staining protocol: cells were firstly stained with a LIVE/DEAD marker (Fixable Blue, UV450, Thermo Fisher Scientific). Secondly, after washing with FACS buffer, anti-mouse TCRγο, ST2, CD11c, CD19, CD103, CD8β, CD45, CD64, CD4, CD11b, MHC-II, CD90.2, and TCR-β were used. After washing, cells were immediately acquired, without fixing, on an LSR Fortessa flow cytometer (BD Biosciences) and analyzed with FlowJo software (Tree Star) using the gating strategy 2 (Supp.Fig.3A-J).

**Table 3:** Anti-mouse antibodies used for immunophenotyping by flow cytometry.

|  |  |  |
| --- | --- | --- |
| Anti-mouse TCR$\gamma\delta$ (clone GL3),<br>PerCP-Cy5.5 | Biolegend | Cat #118118; RRID:<br>AB_10612756 |
| Anti-mouse ST2 (clone RMST2-2), PE | ThermoFisher Scientific<br>(eBioscience) | Cat #25-9335-82;<br>RRID: AB_2637464 |
| Anti-mouse CD11c (clone N418), PE-CF594 | BD Biosciences | Cat #565591; RRID:<br>AB_2869692 |
| Anti-mouse CD19 (clone eBio 1D3),<br>PE-Cy5.5 | ThermoFisher Scientific<br>(eBioscience) | Cat #35-0193-82;<br>RRID: AB_891395 |
| Anti-mouse CD103 (clone 2E7), PE-Cy7 | Biolegend | Cat #121426; RRID:<br>AB_2563691 |
| Anti-mouse CD8β (clone H35-17.2),<br>APC | ThermoFisher Scientific<br>(eBioscience) | Cat #17-0083-81;<br>RRID: AB_657760 |
| Anti-mouse CD45 (clone 30-F11),<br>AF700 | Biolegend | Cat #103128; RRID:<br>AB_493715 |
| Anti-mouse CD64 (clone X54-5/7.1),<br>BV421 | Biolegend | Cat #139309; RRID:<br>AB_2562694 |
| Anti-mouse CD4 (clone GK1.5),<br>BV510 | Biolegend | Cat #100449; RRID:<br>AB_2564587 |
| Anti-mouse CD11b (clone M1/70),<br>BV605 | Biolegend | Cat #101257; RRID:<br>AB_2565431 |
| Anti-mouse MHCII I-A/I-E (clone<br>M5/114.15.2), BV711 | BD Biosciences | Cat #563414; RRID:<br>AB_2738191 |
| Anti-mouse CD90.2 (clone 30-H12),<br>BV780 | Biolegend | Cat #105331; RRID:<br>AB_2562900 |
| Anti-mouse TCRβ (clone H57-597),<br>BUV737 | BD Biosciences | Cat #612821 |
| Anti-mouse IL-17A (clone TC11-18H10.1), PerCP-Cy5.5 | Biolegend | Cat #506920; RRID:<br>AB_961384 |
| Anti-mouse RORγt (clone B2D), PE-CF594 | BD Biosciences | Cat #562684; RRID:<br>AB_2651150 |
| Anti-mouse Tbet (clone 4B10), PE-Cy7 | ThermoFisher Scientific<br>(eBioscience) | Cat #25-5825-82;<br>RRID: AB_11041809 |
| Anti-mouse Gata-3 (clone TWAJ),<br>eFluor660 | ThermoFisher Scientific<br>(eBioscience) | Cat #50-9966-42;<br>RRID: AB_10596663 |
| Anti-mouse IL-13 (clone eBio13A),<br>APC-eFluor780 | ThermoFisher Scientific<br>(eBioscience) | Cat #47-7133-82;<br>RRID: AB_2716964 |
| Anti-mouse FoxP3 (clone FJK-16s),<br>eFluor450 | ThermoFisher Scientific<br>(eBioscience) | Cat #48-5773-82;<br>RRID: AB_1518812 |
| Anti-mouse IFN-γ (clone XMG1.2),<br>BV650 | Biolegend | Cat #505832; RRID:<br>AB_2734492 |
| Anti-mouse IL-10 (clone JES5-16E3),<br>BV711 | BD Biosciences | Cat #564081; RRID:<br>AB_2738581 |

### Luminex and ELISA assays

To evaluate the inflammatory mediators in both small and large intestines, 2.5 cm sections of duodenum, jejunum, ileum, and colon were homogenized for 2 cycles of 20 seconds at 5000 rpm each using Precellys tubes with metal beads in 250 μL of RIPA lysis buffer supplemented with protease inhibitor cocktail (cOmplete^TM^ Mini, Roche). Following centrifugation at 8000 g for 10 minutes at 4°C, the supernatant was used to measure cytokines in each tissue homogenates, using a 32-plex multiplex commercial kit according to the manufacturer’s protocol (Milliplex, Millipore Sigma). IL-25 concentration was determined by ELISA (R&D Systems) and/or Luminex (Millipore Sigma). To determine the antigen-specificity of the cytokine production in the small intestine during *Giardia* infection, 2×10^5^ SI LP cells were cultured in 200uL of supplemented RPMI media alone, or in addition to *Giardia* soluble antigen [10 μg/ml of crude extract from *Giardia intestinalis* trophozoites obtained through sonication cycles on ice], for 24 h in 5% CO_2_ at 37°C. After incubation, the culture supernatant was used for cytokine quantification by a 32-plex Luminex assay (Milliplex, Millipore Sigma) in accordance with the manufacturer’s recommendations.

### Histopathology

For the histopathological analysis, 0.5 cm sections of the proximal duodenum, jejunum, ileum, and colon from *Giardia*-infected mice alone or in the context of *Toxoplasma gondii* co-infection or DSS-induced colitis interaction were fixed in Hollande’s Fixative solution (SatLab) and embedded in paraffin. Five micrometer sections were mounted on glass slides and stained with H&E for inflammatory infiltration analysis, and Periodic Acid–Schiff (PAS) or Alcian Blue Periodic Adic-Schiff (AB/PAS) for mucus production. All images of the tissue sections were scanned using Scan Aperio (Leica).

### Single-cell RNA Sequencing and Data Processing

Live single CD45^+^TCRβ^+^CD4^+^IL10^+^ cells were sorted from isolated lamina propria cell suspension from three pooled *Giardia*-infected mice per sample (total of 4 samples) using a BD FACSAria Cell Sorter (BD Biosciences). Then, approximately 8 × 10^4^ sorted cells (about 2 x 10^4^ per sample) were loaded onto a Chromium Controller (10X Genomics) where individual cells were separated into microdroplets containing beads linked to barcoded oligo-dT. Co-partitioned cells were used in the RNA extraction and library preparation kit Chromium Next GEM Single Cell 3’ Kit v3.1 (10x Genomics) according to the manufacturer’s instructions. Transcripts from lysed cells were reversed transcribed into cDNA and processed by amplification, fragmentation, dual-adapter ligation and size selection. Library quality control was assessed on a 4200 TapeStation (Agilent) and Qubit fluorometer (Life Sciences). Libraries were submitted for high-throughput sequencing on a HiSeqX Ten (Illumina) by Psomagen Inc (Rockville, MD). Sequencing was performed on 4 biological replicates from naïve mice and 4 biological replicates from mice experimentally colonized by *Giardia*.

Illumina raw base calls were demultiplexed into FASTQ files using the mkfastq module in Cell Ranger v.5.0.0 (10x Genomics). FASTQ files were quality-checked using FastQC v.0.11.9 and reads with minimum average quality of 20 and minimum length of 20bp were filtered using fastp v.0.20.1. Cell Ranger v.7.1.0 count was used to align filtered reads to the mouse reference transcriptome mm10 (GENCODE vM23/Ensemble 98) and to quantify the number of different unique molecular identifiers (UMIs) for each gene. The fraction of reads containing a valid barcode assigned to a cell and confidently mapped to the transcriptome varied between 62.6-72.9% in naive and 73.1%-75.7% in infected samples. The estimated number of cells detected in each replicate was between 6,303 and 12,917 cells, for a total of 69,787 cells used as input loaded into RStudio v. 2022.07.0 for downstream processing and analysis using Seurat v.4.3.0.

Cells were filtered-out if mitochondrial gene expression represented > 10% of the total single-cell expression or displayed expression < 800 different genes or > 5000 different genes. Genes detected in fewer than 2 cells were removed. UMI counts were normalized by library read depth, log-transformed, centered and Z-scored using the functions NormalizeData and ScaleData. Highly variable genes were identified using FindVariableFeatures (2000 features, “vst”method). Data integration was performed by selecting genes that are variable in most samples with SelectIntegrationFeatures and identifying cells to be used as “anchors” with FindIntegrationAnchors, to integrate the different samples using the IntegrateData function. Dimensionality reduction was performed on scaled integrated data using RunPCA and RunUMAP and clustering with FindNeighbors and FindClusters with 27 principal components and a resolution of 0.2. Transcript markers for each cell cluster were identified using the FindAllMarkers function and evaluating the average expression and the percent expressed of each gene in a cluster compared to others. Markers were selected if single-cell log2FC expression > 0.25 and statistically significant (Wilcoxon rank-sum test adjusted p-value < 0.05). Clusters were manually annotated according to the top 10 gene markers and reference-based analysis using SingleR v.1.10.0 to compare each cluster gene expression with mouse bulk RNA-seq data available on the Immunological Genome Project (Immgen). Pathway enrichment analysis was performed using the top markers in Th2 cells (avg log2FC>0.25) as input for clusterProfiler v.4.8.2 and enrichplot v1.20.0 in R with a Benjamini-Hochberg q-value<0.05. The scRNA-seq raw data files generated in this study were deposited in the NCBI BioProject database under accession code PRJNA1082359.

### Recombinant IL-10, IL4, and IL-13 treatment

Naïve mice were intraperitoneally treated with 250 ng of recombinant mouse IL-10 (Peprotech), and/or with 10 μg of both combined recombinant mouse IL-4-Fc and IL-13-Fc (5 μg of each) (Absolute Antibody) every other day for 10 days (5 treatments), before and during *Toxoplasma gondii* infection. Intraperitoneal injections of PBS were used as a control.

### In vivo Imaging

*Toxoplasma gondii*-infected mice were imaged for bioluminescent detection of firefly luciferase activity seven days post-infection using an IVIS BLI system from Xenogen to monitor parasite burden. Five to ten minutes before imaging, the infected mice were injected with 3 milligrams (diluted in 200 μl of PBS) of D-luciferin (PerkinElmer) substrate, and then they were imaged for 300 seconds to detect emitted photons.

### Statistical Analysis

Scatter plots and statistical analyses were performed using Prism 9.0 software (GraphPad). For groups of two, the test for significance was conducted using the non-parametric Mann-Whitney test, and for groups of three or more, one-way analysis of variance (ANOVA) followed by Sidak’s multiple comparisons test was applied to assess significance. Differences were considered significant when the *p* value was ≤ 0.05.

## ACKOWLEDGMENTS

We are grateful to Dr. Balogun Joshua, Dr. Dahal Samuel, Mr. Muhammad Ali, Dr. Njom Victor, Dr. Amana Onekutu, and Dr. Emmanuel Effanga who coordinated the samples collection in Nigeria, as well as administered the questionnaire used to establish whether infection was associated with symptomatic disease. We also thank Dr. Camila de Oliveira Silva Souza and Dr. P’ng Loke for providing us the STAT6^-/-^ mice and the mouse recombinant IL-4-Fc and IL-13-Fc cytokines. This work was supported by the Division of Intramural Research of the National Institute of Allergy and Infectious Diseases (NIAID) at the National Institutes of Health, and NIH extramural grant AI109591 to SMS.

## AUTHOR CONTRIBUTIONS

Conceptualization, A.S.-S., C.H.C., S.M.S., and M.E.G.; Methodology, A.S.-S., P.H.G.-G., C.H.C., and T.R.F.; Investigation, A.S.-S., P.H.G.G., O.G.A., T.R.F., E.V.C.A.-F., E.T.T., B.G., and M.Y.F.; Writing – Original Draft, A.S.-S. and P.H.G.G.; Writing – Review & Editing, A.S.-S., P.H.G.-G., S.M.S., M.E.G.; Funding Acquisition, S.M.S. and M.E.G.; Resources, S.M.S. and M.E.G.; Supervision, M.E.G.

## SUPPLEMENTAL FIGURES AND LEGENDS

**Supplemental figure 1:**
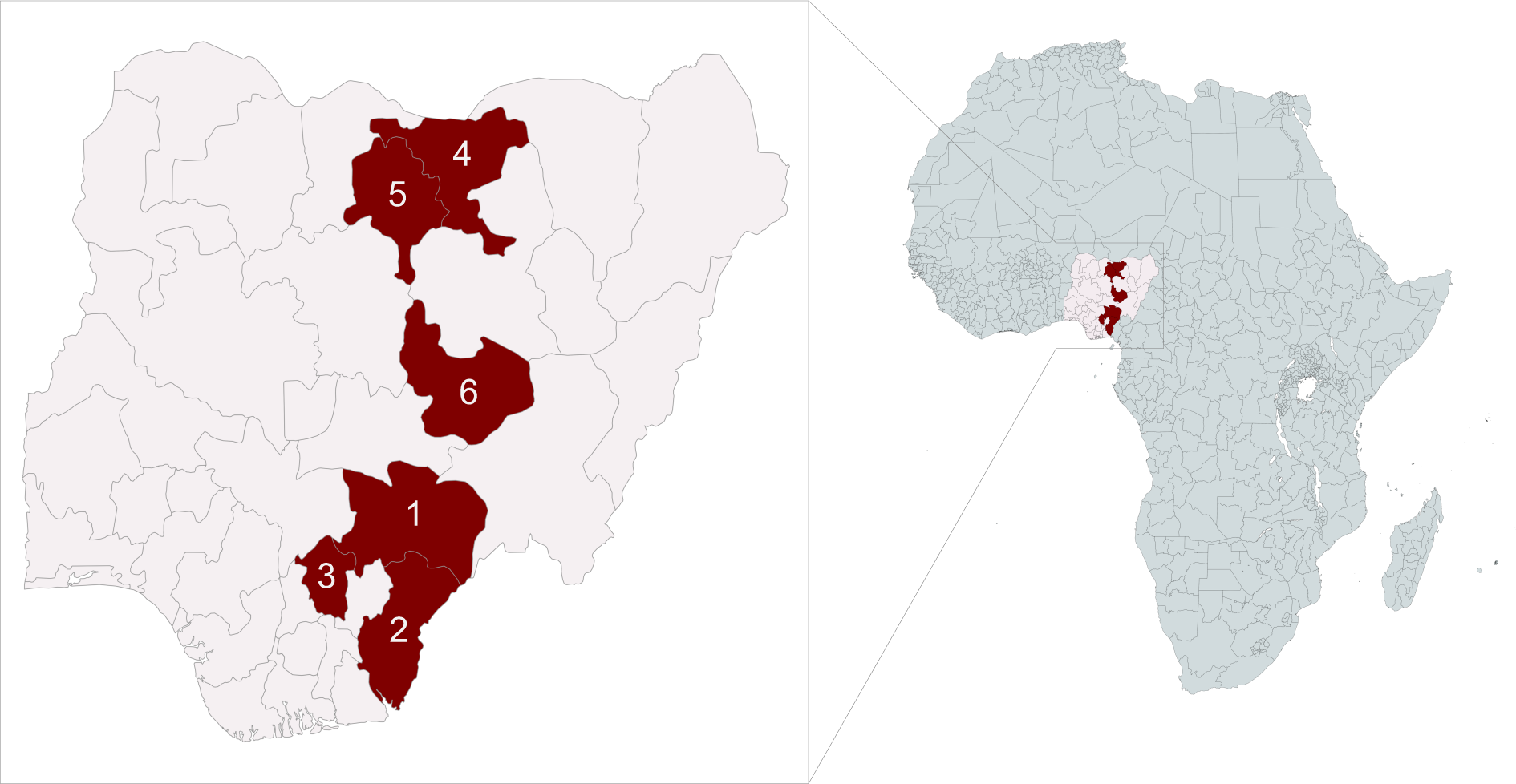
Nigeria map showing sampled states. 1. Benue; 2. Cross River; 3. Enugu; 4. Jigawa; 5. Kano; 6. Plateau.

**Supplemental figure 2:**
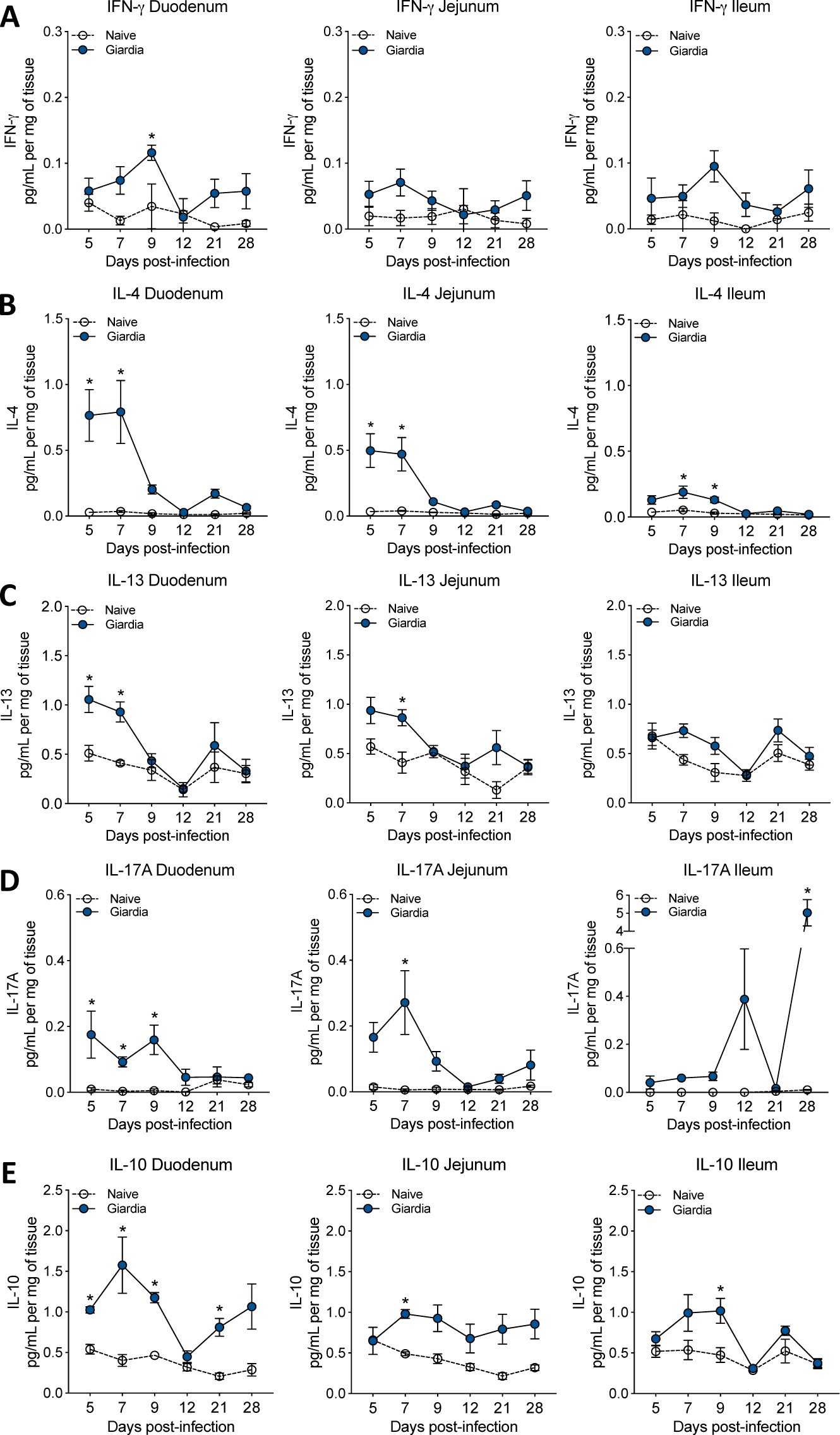
Cytokine production kinetics across the three segments from the small intestine in *Giardia*-infected mice. Levels of IFN-γ (A), IL-4 (B), IL-13 (C), IL-17A (D), and IL-10 (E) in the three different sections of the small intestine tissue homogenate (normalized by mg of tissue) from naïve or *Giardia*-infected mice at 5-, 7-, 9-, 12-, 21-, and 28-days post-infection (measured by Luminex). Data are represented as mean ± SEM for each time point and significance was calculated with one-way ANOVA test followed by Sidak’s multiple comparisons test. *p≤0.05, **p≤0.01, ***p≤0.001. Data are representative of two independent experiments.

**Supplemental figure 3:**
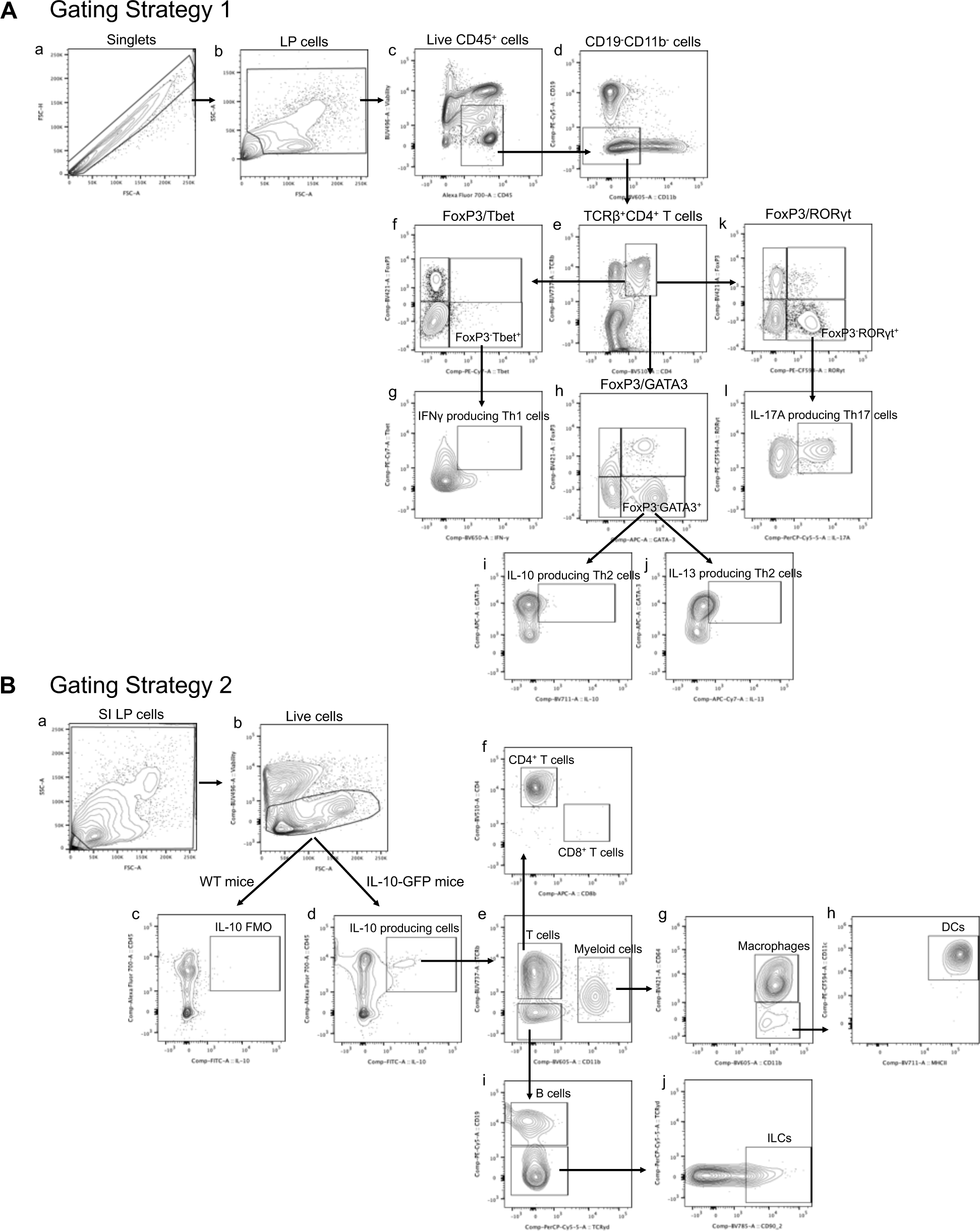
Gating strategy for immunophenotypic analysis by Flow Cytometry. (A) The gating strategy 1 was used for the overall immunophenotypic analysis of the experiments reported in figures 1B, 1G, 2D, 2G, 4B, 4C, 5E, 5J, 6E, and 6J. Briefly, singlets (a) were gated followed by subsequent lamina propria cell gating by FSC-A vs SSC-A (b). (c) Live hematopoietic cells were gated as CD45/Alexa Fluor700^+^ and Live/Dead/UV496^-^. (d) B cells and myeloid cells were then excluded by gating as CD19/PE-Cy5^-^ and CD11b/BV605^-^. (e) CD4^+^ T cells were further gated as TCRβ/BUV737^+^ and CD4/BV510^+^. Subsets of CD4^+^ T cells were further subdivided into either the single or co-expression of FoxP3/BV421 in combination with single or co-expression of Tbet/PE-Cy7 (f), GATA3/APC (h) or RORγt/PE-CF594 (k). Finally, IFN-γ/BV650 (g), IL-10/BV711 (i), IL-13/PE (j), and IL-17A/PerCP-Cy5.5 (l) were analyzed into the populations of Tbet^+^ Th1 cells, GATA3^+^ Th2 cells, and RORγt^+^ Th17 cells, respectively. **(B)** The gating strategy 2 was design specifically for the identification of the major sources of IL-10 in the lamina propria of either naïve-and *Giardia*-infected IL-10 GFP reporter mice (Figure 2F). For this, live CD45/Alex Fluor 700^+^ lamina propria cells (a, b) were gated for the IL-10 GFP expression (d), using a WT mouse as a control for a negative gate (c). (e) IL-10 producing hematopoietic cells were further characterized as either TCRβ/BUV737^+^ T cells, or CD11b/BV605^+^ cells or TCR3^-^CD11b^-^ cells. (f) T cells were subdivided as CD4/BV510^+^ cells or CD8β/APC^+^ cells. (g) CD64/BV421^+^ macrophages were identified within the CD11b^+^ cells; and (h) CD64/BV421^-^, CD11c^+^/PE-CF596+, MHCII^+^/BV711^+^ cells were classified as Dendritic Cells (DCs). (i) Non-T and non-myeloid cells were further gated as CD19/PE-Cy5^+^ cells or TCRγο/PerCP.Cy5.5 T cells. (j) Finally, IL-10 producing CD45^+^TCR3^-^CD11b^-^CD19^-^TCRγο^-^ CD90/BV785^+^ cells were classified as innate lymphoid cells (ILCs).

**Supplemental figure 4:**
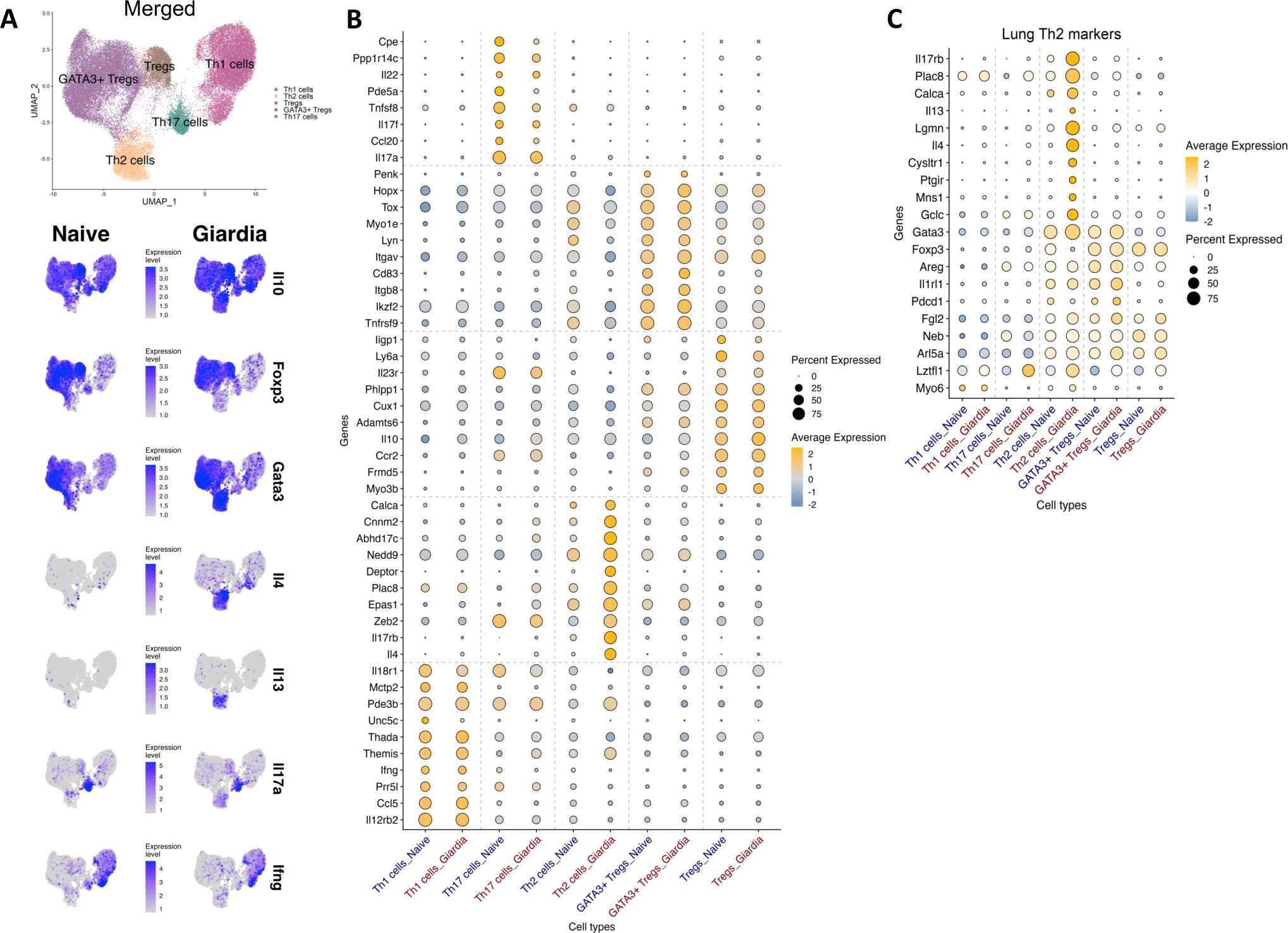
Molecular characterization of IL-10 producing CD4^+^ T cells in the small intestine lamina propria. (A) UMAP plot showing the clustering analysis of the merged dataset of all sorted IL-10 producing CD4^+^ T subsets in the small intestine lamina propria from both naïve and *Giardia*-infected IL-10-GFP reporter mice. The plot distinctly identifies five key clusters, corresponding to Th1, Th2, Th17, Treg, and Th2 Treg cells subset; and feature plot showing the gene expression level of *Il10*, *Foxp3* and *Gata3, Il4, Il13, Il17a, Ifng* among the major clusters of each IL-10 producing CD4^+^ T subset from naïve (left) or *Giardia*-infected mice (right). (B) Dot plot graph highlighting the top10 highly expressed genes within each cluster of IL-10 producing CD4^+^ T cells in the small intestine lamina propria from naïve and *Giardia*-infected mice. (C) Dot plot graph highlighting the signature genes of pathogenic Th2 cells within each cluster of IL-10 producing CD4^+^ T cells in the small intestine lamina propria from naïve and *Giardia*-infected mice.

**Supplemental figure 5:**
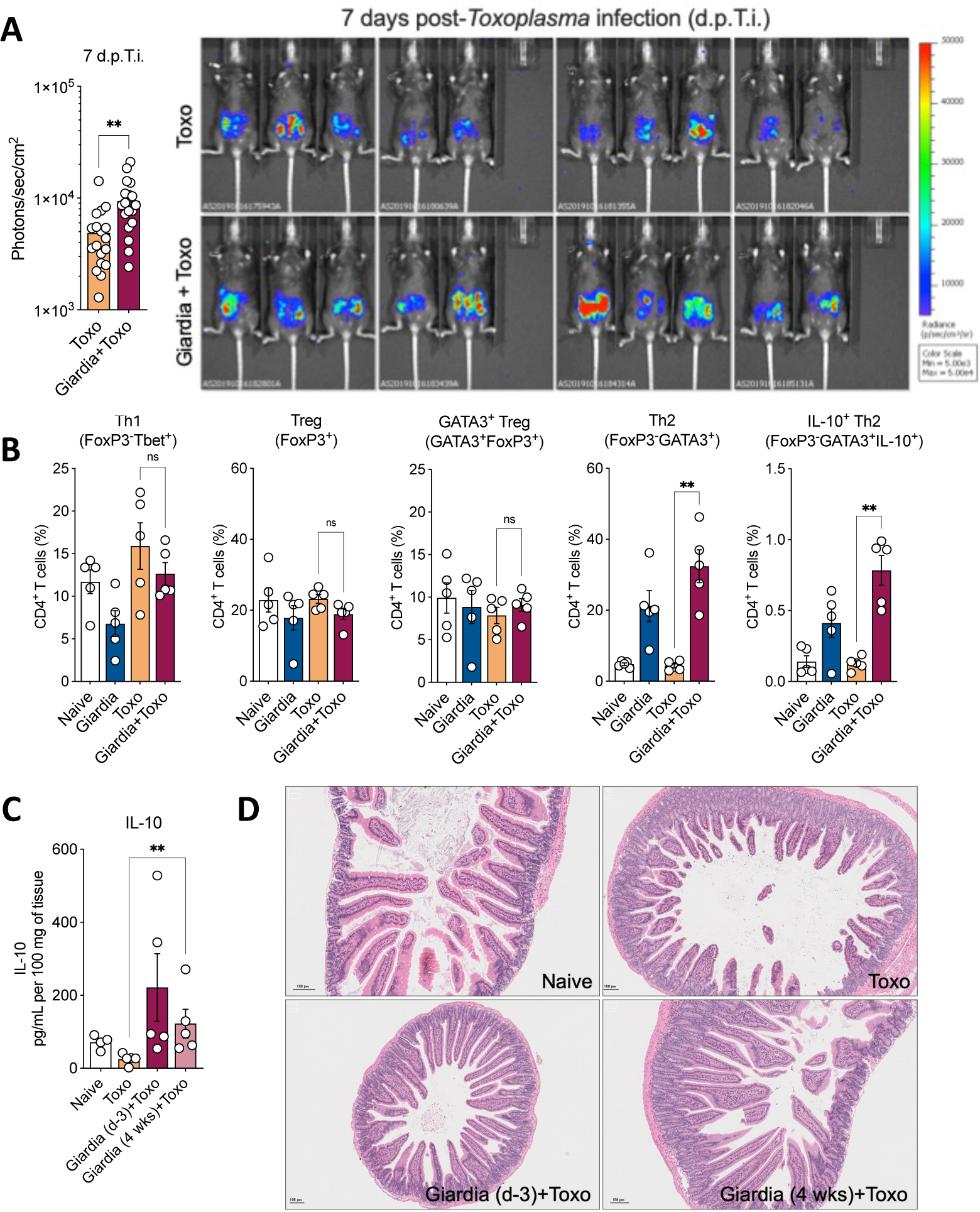
The effect of *Giardia* chronicity in the co-infection with *T. gondii*. (A) Bioluminescent detection in photons/sec/cm^2^ shows *Toxoplasma* burden *in vivo* (7 d.p.T.i.) in mice co-infected or not with *Giardia* GS/M strain three days before. (B) Scatter plot graphs indicating the frequency of Th1 (FoxP3^-^Tbet^+^), total Treg (FoxP3^+^), GATA3^+^ Treg (GATA3^+^FoxP3^+^), Th2 (GATA3^-^Foxp3^+^), and IL-10^+^ Th2 (FoxP3^-^GATA3^+^IL-10^+^) cells in the small intestine lamina propria of *Giardia*-(three days before, d-3), *Toxoplasma*-, or *Giardia* (d-3)+*Toxoplasma*-infected mice 4 days post-*Toxoplasma* infection (4 d.p.T.i.). Gated on Live CD45^+^TCR3^+^CD4^+^. (C) IL-10 levels in the proximal small intestine of *Giardia*-chronically infected mice (4 weeks post-infection, 4 wks). (D) Representative image of H&E staining of the proximal small intestine from naïve, *Toxoplasma-*, *Giardia* (d-3)+*Toxoplasma*-, or *Giardia* (4 wks)*+Toxoplasma*-infected mice (8 days-post *Toxoplasma* infection). Scale bars represent 100 μm. Data are represented as mean ± SEM for each time point and significance was calculated with one-way ANOVA test followed by Sidak’s multiple comparisons test. *p≤0.05, **p≤0.01, ***p≤0.001. Data are representative of two independent (A-B) and one (C-D) experiment.

